# A CITE-seq atlas of mouse embryonic hematopoiesis maps endothelial heterogeneity and reveals emerging HSC-associated states

**DOI:** 10.64898/2026.11.17.767285

**Authors:** Daniel García Sánchez, Vrinda Sharma, Alberto González González, Carolina Sañudo Campo, Katie Stoker, Sudhakar Sahoo, Martin Proffitt, Michael Lie-A-Ling, Flor Perez-Campo, Georges Lacaud, Roshana Thambyrajah

## Abstract

The emergence of hematopoietic stem cells (HSCs) from hemogenic endothelium is a rare developmental event, yet how transcriptional identities relate to cell-surface phenotypes during this process remains poorly understood. Here, we generated a CITE-seq atlas of mouse embryonic hematopoiesis spanning E9.5, E10.25 and E11.5 aorta-gonad-mesonephros (AGM) regions and E11.5 foetal liver. Integrated single-cell transcriptomic and surface-protein profiling revealed extensive endothelial and hematopoietic heterogeneity, distinguishing broad hemogenic specification from more restricted HSC-associated transcriptional states. Notably, *Hlf* expression closely aligned with HSC-associated gene programmes and CD27 surface-protein abundance, identifying a potential marker for the prospective enrichment of HSC-associated populations.

Graph-based connectivity, diffusion pseudotime and RNA velocity analyses resolved distinct arterial preHSC-HE and HSC-HE populations and supported an inferred progression towards emerging HSC-associated states. Strikingly, we identified a rare subset of E9.5 endothelial cells (12 of 101) exhibiting elevated predicted connectivity towards arterial preHSC-HE without detectable *Runx1* expression. These cells displayed distinct transcriptional and cell-surface profiles, suggesting that molecular heterogeneity associated with hemogenic specification may arise before the activation of established hemogenic programmes. Furthermore, the inferred progression towards HSC-associated identity was accompanied by coordinated changes in gene expression and surface-protein abundance, revealing phenotypic features associated with successive developmental states.

Together, our findings uncover previously unresolved endothelial heterogeneity associated with early HSC development and establish a link between emerging HSC-associated transcriptional identities and their cell-surface phenotypes. This multimodal atlas provides a framework for the prospective isolation and functional characterization of rare endothelial populations potentially involved in HSC specification.

## Introduction

In the mouse embryo, definitive HSCs emerge during mid-gestation in the aorta-gonad-mesonephros (AGM) region through endothelial-to-hematopoietic transition (EHT), a remarkable process in which specialized hemogenic endothelial (HE) cells progressively relinquish their endothelial identity to acquire a hematopoietic fate ^1–5^. Yet, the emergence of hematopoietic cells does not necessarily mark the birth of an HSC. Within the same developmental environment, a much larger and heterogeneous population of hematopoietic progenitors (HPCs) is generated, contributing to the establishment of the embryonic blood system without acquiring the lifelong regenerative and multi-lineage potential of HSCs ^6–9^. These emerging populations form intra-aortic hematopoietic clusters (IAHCs) ^4, 10^, whose cellular and functional heterogeneity remains incompletely understood ^11,12^. Despite decades of investigation, no universally consistent combination of surface markers reliably distinguishes functional HSCs from closely related progenitors within the AGM, and a comprehensive immunophenotypic characterization of the cellular diversity within IAHCs remains lacking. Consequently, although the transition from endothelium to blood has been extensively characterized ^11–15^, our ability to distinguish the rare cells acquiring lifelong stem cell potential from the multitude of emerging hematopoietic progenitors remains remarkably limited. A particularly important challenge is to distinguish cells that have initiated an HSC-associated transcriptional programme from the broader population of hematopoietic precursors. The transcription factor *Hlf* (Hepatic leukemia factor) has emerged as a prominent molecular feature of developing hematopoietic stem and progenitor populations ^16^ ^17^ ^7^ ^18^. However, lineage-tracing studies have demonstrated that HLF-expressing embryonic precursors can contribute to both HSC and progenitor populations, indicating that HLF expression alone does not define an exclusively stem cell-restricted fate ^7^. Moreover, although combinations of surface markers have improved the prospective identification of emerging HSCs, a systematic comparison of the cellular populations identified by individual markers and their combinations remains lacking, leaving their relationship to distinct transcriptional and developmental identities poorly understood. ^19^.

Single-cell transcriptomic technologies have provided unprecedented opportunities to investigate these developmental transitions, revealing cellular heterogeneity that is often concealed within conventionally defined populations ^15,16,20^. However, transcriptional identity alone provides an incomplete description of cellular state, as transcript and protein abundance can reflect distinct aspects and timescales of cellular identity. Integrating these complementary molecular layers may therefore provide a more comprehensive view of developmental heterogeneity and the transitions between cellular states ^21,22^.

Here, we combine single-cell transcriptomic profiling with antibody-derived surface protein measurements by CITE-seq (Cellular Indexing of Transcriptomes and Epitopes by sequencing) of an Immune panel, in combination with endothelial, HSC-associated and Notch signaling surface markers to investigate the molecular and phenotypic diversification of hematopoietic populations during mouse embryonic development. By examining successive developmental stages spanning early embryonic hematopoiesis, the E9.5, E10.25 and E11.5 AGM and E11.5 foetal liver, we resolve distinct endothelial, hemogenic and hematopoietic populations and investigate their transcriptional relationships using RNA velocity and pseudotemporal analyses. We further integrate these molecular states with surface protein expression to examine how HSC-associated transcriptional identity relates to phenotype. Together, our study provides a framework for understanding how HSC-associated molecular identity emerges within the developing hematopoietic system and how transcriptional and phenotypic information may help distinguish rare stem cell-associated populations from their closely related progenitors.

## Results

### Single-cell multiomic profiling reveals extensive molecular and phenotypic heterogeneity during embryonic haematopoiesis

We performed single-cell RNA sequencing (scRNA-seq) coupled with antibody-derived tag (ADT) profiling ^21,23^ of cells isolated from the mouse AGM region at E9.5 (n=9), E10.25 (n=23) and E11.5 (n=10), together with E11.5 foetal liver, to investigate the transcriptional and cell-surface phenotypic heterogeneity associated with emerging hematopoietic populations (Fig. 1A, B). Single cell suspension from each sample was simultaneously stained with a panel of Totalseq B ADTs (suppl Table S1), and fluorescently labelled CD31 and cKIT.

**Figure 1.**
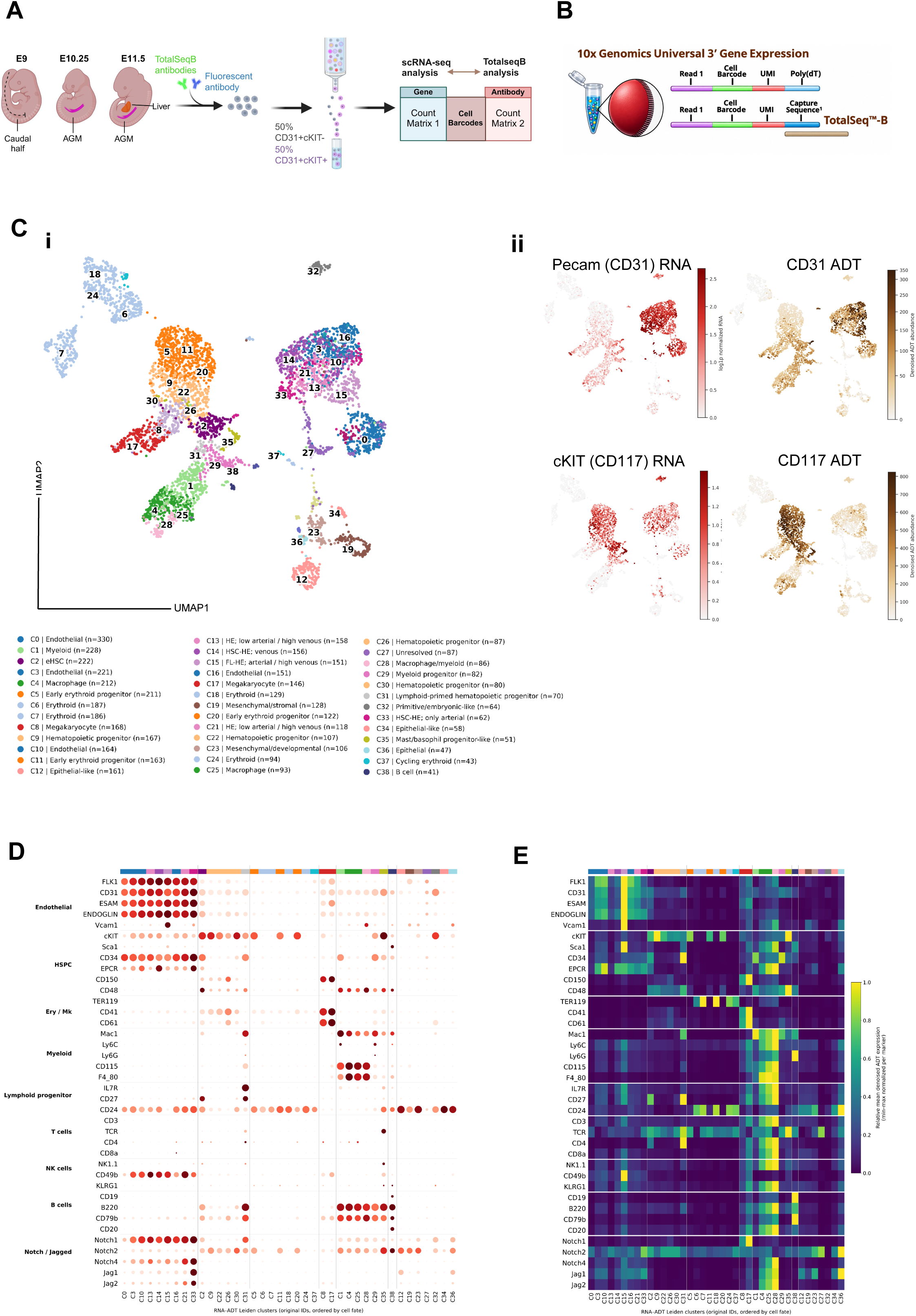
Single-cell multimodal profiling reveals transcriptional and cell-surface heterogeneity during embryonic hematopoiesis. **(A)** Schematic of the experimental workflow. Cells were isolated from mouse embryonic tissues at E9.5, E10.25 and E11.5, including the aorta-gonad-mesonephros (AGM) region and E11.5 foetal liver. Following enrichment based on CD31 and c-KIT expression, cells were stained with TotalSeq-B antibodies and processed for simultaneous single-cell RNA and surface-protein profiling. **(B)** Schematic of the 10x Genomics gene-expression and antibody-derived tag (ADT) capture workflow**. (C)** Integrated single-cell atlas generated using totalVI. **(i)** Uniform manifold approximation and projection (UMAP) showing 39 annotated cell clusters, encompassing endothelial, hemogenic endothelial and hematopoietic populations**. (ii)** UMAP feature plots comparing transcript expression and ADT-derived protein abundance for CD31 (*Pecam1*) and c-KIT (*Kit*; CD117). **(D)** Dot plot showing expression of selected lineage-associated genes across the annotated cell clusters. Dot size indicates the proportion of cells expressing each gene, and colour represents relative expression abundance. **(E)** Heatmap showing ADT-derived surface-protein abundance across the annotated clusters. Markers are grouped according to endothelial, hematopoietic, erythroid, myeloid, lymphoid and NOTCH-associated categories. Colour intensity indicates relative protein abundance.

Cells were enriched using CD31 and c-KIT through flow cytometric sorting, allowing us to capture endothelial (CD31+cKIT-), hemogenic (CD31+cKIT-) and emerging hematopoietic populations (CD31+cKIT+) across successive developmental stages at a 50%/50% ratio (Fig 1A and Suppl Fig 1A). Subsequently, the purified cells were loaded onto 10x genomics universal 3’ gene expression ChIP (Fig 1B) and run through the GEM-X OCM workflow. The samples were sequenced, and the sequencing reads for gene expression and the separate library for ADT counts were processed, quality controlled and derived from the dedicated 10x genomics CellRanger application. Since gene expression and surface protein abundance were measured simultaneously using the 10x Genomics platform and TotalSeq-B ADT antibodies, we obtained complementary transcriptional and immunophenotypic information for individual cells.

First, we used unsupervised clustering using UMAP (Uniform Manifold Approximation and Projection, ^24^) of the integrated transcriptomic and ADT (totalVI ^25^ denoised) dataset labelled 39 transcriptionally distinct populations. To assign cell identities to each cluster, we used differentially expressed genes (DEGs) across Leiden clusters together with ADT-based immunophenotyping to annotate all 39 populations in our dataset (Fig. 1Ci and Supplementary Table1). The fully annotated UMAP encompasses a diverse range of endothelial and hematopoietic populations in the AGM, as well as HSPCs present in the E11.5 foetal liver (Fig. 1Ci and Supplementary Fig. 1B). To assess transcript–protein correspondence, we compared *Pecam1*/CD31 and *Kit*/CD117 expression, as these markers were used for cell enrichment (Fig. 1A). As expected, *Pecam1*/CD31 was enriched in endothelial clusters, whereas *Kit/*CD117 was enriched in hematopoietic clusters (Fig. 1Cii). Importantly, transcript expression and ADT-derived protein abundance showed broadly corresponding distributions across the annotated populations (Fig. 1Cii). We further examined established lineage-associated genes distinguishing endothelial, hematopoietic stem and progenitor, erythroid, megakaryocytic, myeloid and lymphoid populations, together with NOTCH signalling components, and compared their expression patterns with the corresponding surface-protein profiles (Fig. 1D,E). These analyses revealed consistent lineage-associated transcriptional and immunophenotypic patterns across the annotated clusters, supporting the reliability of the ADT-derived protein measurements.

Together, these analyses demonstrate the utility of CITE-seq for characterizing the transcriptional and immunophenotypic landscape of early hematopoietic development at the single-cell level.

### Developmental progression is accompanied by profound changes in hematopoietic cell composition

We next investigated how the cellular composition of the hematopoietic compartment changes during embryonic development. To further characterize the transcriptionally defined populations of interest, we examined the expression of gene signatures associated with endothelial, arterial, venous and hematopoietic identities, including programmes characteristic of hemogenic endothelium (HE) and hematopoietic stem cells (HSCs) (Fig. 2A,B and Supplementary Fig. 2B,C). UMAP visualization of these gene programmes allowed us to identify clusters enriched for each transcriptional signature (Fig. 2B). The endothelial programme, comprising *Cdh5*, *Kdr*, *Pecam1*, *Esam* and *Eng*, was broadly enriched across the endothelial compartment (Fig. 2B). In contrast, arterial and venous programmes displayed more restricted distributions, identifying distinct populations within the broader endothelial landscape. The arterial signature, including *Efnb2*, *Gja5*, *Dll4*, *Hey2* and *Sox17*, was enriched within a subset of endothelial clusters, whereas the venous programme, including *Ephb4*, *Nr2f2* and *Nrp2*, showed a largely distinct distribution (Fig. 2B). These differences were further supported by the developmental-stage-specific programme scores, which showed that endothelial identity was broadly maintained across endothelial populations, while arterial and venous signatures remained distinguishable across the analysed stages (Supplementary Fig. 2C).

**Figure 2.**
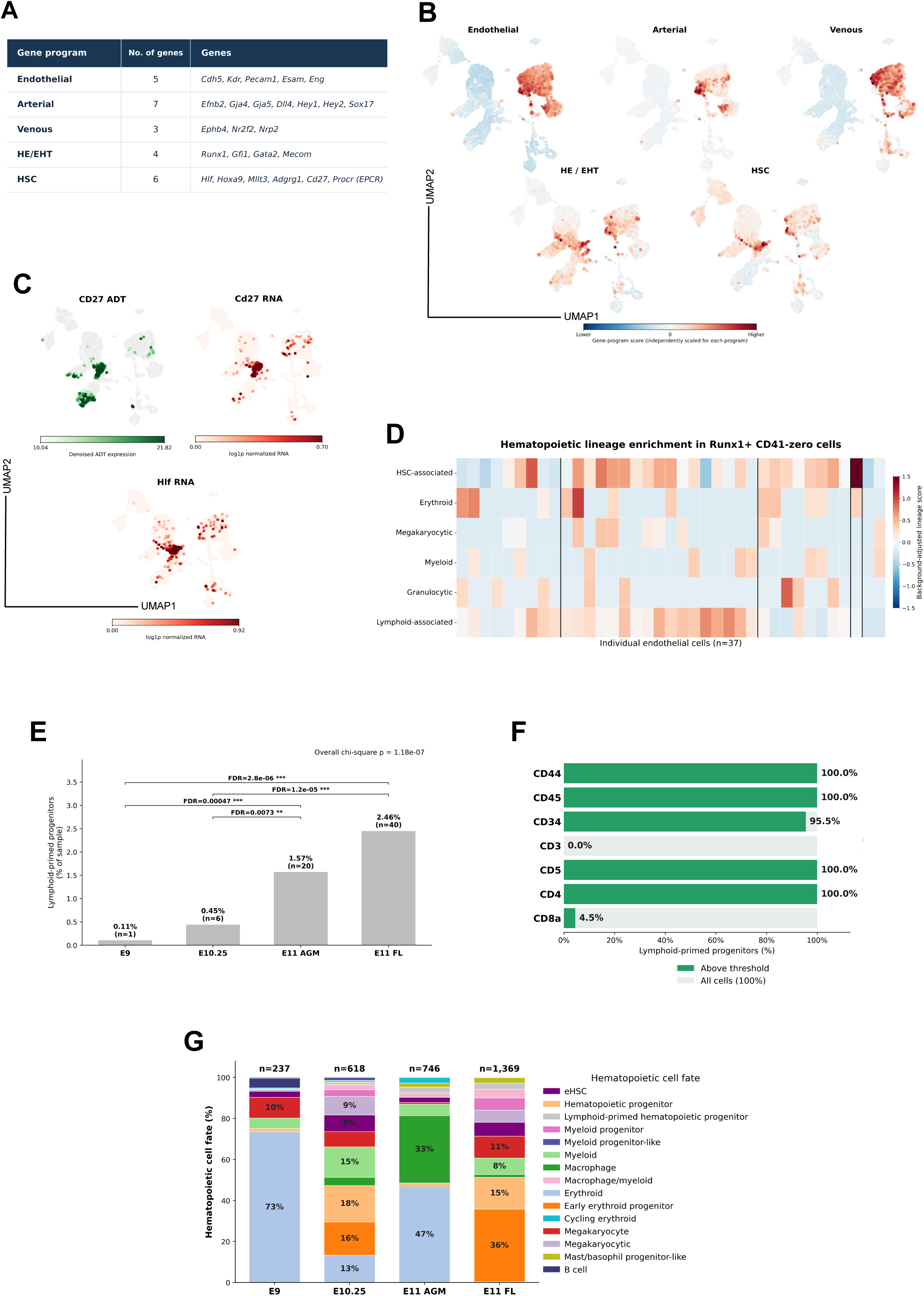
Multimodal characterization identifies distinct endothelial and hematopoietic populations during embryonic development. **(A)** Gene sets used to calculate transcriptional programme scores associated with endothelial, arterial, venous, hemogenic endothelial/endothelial-to-hematopoietic transition (HE/EHT) and hematopoietic stem cell (HSC) identities. **(B)** UMAP feature plots showing the distribution of endothelial, arterial, venous, HE/EHT and HSC-associated transcriptional programme scores across the integrated dataset. Colour intensity indicates relative programme enrichment. **(C)** UMAP feature plots comparing CD27 surface-protein abundance, *Cd27* transcript expression and *Hlf* transcript expression. **(D)** Heatmap showing enrichment of HSC-associated, erythroid, megakaryocytic, myeloid, granulocytic and lymphoid-associated transcriptional programmes among individual *Runx1*-expressing endothelial cells lacking detectable CD41 surface protein (n = 37). **(E)** Frequency of transcriptionally defined lymphoid-primed progenitors across E9.5, E10.25, E11.5 AGM and E11.5 foetal liver samples. Bars indicate the percentage of cells assigned to this population, with cell counts shown above each bar. Statistical comparisons and false discovery rate (FDR)-adjusted values are indicated. **(F)** Percentage of lymphoid-primed progenitors expressing selected surface markers above the applied ADT threshold, including CD44, CD45, CD34, CD3, CD5, CD4 and CD8a. **(G)** Relative composition of the captured hematopoietic compartment across developmental stages. Stacked bars indicate the proportion of transcriptionally annotated hematopoietic populations within each sample, with total cell counts shown above the bars.

The HE/EHT programme, defined by *Runx1*, *Gfi1*, *Gata2* and *Mecom*, exhibited a different pattern of enrichment (Fig. 2A,B and Supplementary Fig. 2B,C). Rather than being restricted to a single endothelial population, this signature extended across transcriptionally related populations within the endothelial-to-hematopoietic landscape. This distribution is consistent with the activation of hematopoietic programmes during endothelial-to-hematopoietic transition (EHT) and suggests that hemogenic specification encompasses multiple molecular states. Interestingly, the HE/EHT programme partially overlapped with the HSC-associated signature, comprising *Hlf*, *Hoxa9*, *Mllt3*, *Adgrg1*, *Cd27* and *Procr* (Fig. 2A,B). However, the two programmes did not completely coincide. While the HE/EHT signature marked a broader distribution of cells associated with hematopoietic specification, the HSC-associated programme was enriched within a more restricted region of the UMAP (Fig. 2B). This distinction was also evident in the programme-score heatmaps, which revealed differences in HE/EHT and HSC-associated signatures across endothelial and hemogenic populations and developmental stages (Supplementary Fig. 2C).

These programme scores further supported the annotation of distinct endothelial and HSC-associated populations. Notably, we identified a population of arterial endothelial cells displaying low venous programme scores but elevated HE/EHT and HSC-associated scores, particularly evident in the developmental-stage-specific analysis (Supplementary Fig. 2C). We designated this C33 cluster as arterial HSC-HE. In contrast, the HSC-associated programme was strongly enriched in cluster C2, which contained a substantial representation of cells from the E11.5 AGM and foetal liver, consistent with the developmental emergence of HSCs and their subsequent colonization of the foetal liver (Fig. 2B and Supplementary Fig. 2C). We further characterized this emerging HSC-associated (eHSC) cluster by examining the expression of established HSC-associated markers, including *Hlf*, *Cd27*, *Hoxa9*, *Procr*, *Runx1* and *Mllt3* (Fig. 2B,C and Supplementary Fig. 2A,B). These markers showed overlapping enrichment within the C2/eHSC cluster. Moreover, CD27 ADT abundance was strongly enriched in this population (Fig. 2C), supporting the association between its transcriptional identity and surface-protein phenotype.

Given the diversity and heterogeneity observed among endothelial and HE populations, we next investigated whether lineage-associated transcriptional programmes could reveal further heterogeneity within the early hemogenic compartment. To this end, we pooled the endothelial clusters and examined CD41 ADT abundance, as CD41 is an early surface marker associated with EHT (Supplementary Fig. 3A). Most endothelial cells lacked detectable CD41 ADT. We then further classified these cells according to *Runx1* expression, a key regulator of hemogenic specification^26,27^. Specifically, we selected endothelial cells lacking detectable CD41 ADT but expressing *Runx1*, representing a candidate HE-enriched population, and examined their hematopoietic lineage-associated gene programme scores (Fig. 2D and Supplementary Fig. 3B). The resulting heatmap revealed considerable transcriptional heterogeneity within this early HE-associated compartment, with cells displaying differential enrichment of HSC-, erythroid-, megakaryocytic-, myeloid-, granulocytic- and lymphoid-associated programmes. Interestingly, lymphoid-associated and HSC-associated programmes showed partial overlap (Fig. 2D). The relative representation of lymphoid-primed progenitors increased across the developmental samples, from approximately 0.11% at E9.5 to 0.45% at E10.25, 1.57% in the E11.5 AGM and 2.46% in the E11.5 foetal liver (Fig. 2E). Thus, lymphoid-associated transcriptional programmes were detectable during early embryonic hematopoiesis and became progressively more represented at later developmental stages.

Next, to investigate the surface-protein characteristics of this population, we examined the ADT abundance of selected markers. Lymphoid-primed progenitors displayed widespread detection of CD44, CD45, CD4 and CD8a, together with CD34 in most cells, whereas CD3 and CD8b were detected in only a small fraction of cells or remained undetectable according to the applied expression threshold (Fig. 2F). These findings revealed a distinctive combination of surface-protein expression within a transcriptionally defined lymphoid-primed progenitor population, illustrating how paired transcriptomic and immunophenotypic measurements can resolve heterogeneity among developing hematopoietic populations.

To obtain a broader overview of hematopoietic cell types and their relative abundance within the captured intra-aortic hematopoietic cluster (IAHC)-associated compartment, we next assessed hematopoietic cell composition across developmental stages. At E9.5, the captured CD31⁺c-KIT⁺ hematopoietic compartment was dominated by erythroid cells, which accounted for approximately 73.4% of the analysed cells. By E10.25, the relative contribution of erythroid cells was markedly reduced, accompanied by an increased representation of myeloid and other hematopoietic progenitor populations (Fig. 2G and Supplementary Fig. 1B). At E11.5 in the AGM, erythroid cells remained abundant, representing approximately 46.9% of the captured hematopoietic cells, while myeloid populations also constituted a substantial fraction. In contrast, the E11.5 foetal liver displayed a distinct cellular composition, with a greater representation of erythroid progenitors and differentiating hematopoietic populations (Fig. 2G and Supplementary Fig. 1B). Together, these findings reveal heterogeneity in lineage-associated transcriptional programmes within the early endothelial compartment, alongside pronounced developmental and anatomical differences in the composition of the sampled hematopoietic populations.

### Transcriptional connectivity and pseudo temporal analyses resolve distinct endothelial and HSC-associated states

Having established the molecular diversity of the developing hematopoietic compartment, we subsequently sought to investigate the relationships between endothelial, hemogenic and HSC-associated populations. Although HSCs arise through EHT, the extent to which emerging HSC-associated states represent a continuous developmental progression or distinct transcriptional populations remains incompletely understood.

To investigate the developmental relationships between endothelial and HSC-associated populations, we used complementary bioinformatical tool. We assessed transcriptional connectivity between clusters using Partition-based Graph Abstraction (PAGA^28^). In parallel, we also implemented RNA velocity driven probabilities^29,30^. Whereas PAGA assesses connectivity between cell populations based on the structure of the single-cell nearest-neighbor graph^28^, RNA velocity estimates the direction of transcriptional change by modeling the relative abundances and kinetics of spliced and unspliced transcripts ^29,30^. These velocity-derived transition probabilities can subsequently be used to assess the relative likelihood of transitions between transcriptionally defined cellular states and, when combined with complementary downstream analyses, infer their relative positioning along putative developmental trajectories. Although graph-based connectivity and RNA velocity use different computational approaches, both provide complementary information about potential relationships between cellular states. By quantifying connectivity between cell clusters within the single-cell nearest-neighbour graph, partition-based graph abstraction (PAGA) revealed a network linking endothelial and hemogenic populations (Fig. 3A), consistent with potential developmental relationships during early hematopoiesis. Notably, the HSC-associated cluster C2 occupied a more restricted position within this connectivity network and was linked to the endothelial compartment through cluster C33, annotated as arterial HSC-HE (Fig. 3A). This observation suggested that not all endothelial populations shared equally strong graph-based relationships with C2/eHSC. We therefore examined the developmental-stage representation of the endothelial and HSC-associated populations identified in this analysis (Supplementary Fig. 3C). Arterial preHSC-HE cells were predominantly represented at E9.5, whereas other endothelial and HE-associated populations showed greater contributions from E10.25 and E11.5 AGM. In contrast, the eHSC population was more strongly represented in the E11.5 foetal liver, consistent with developmental and anatomical differences in the distribution of these transcriptional states (Supplementary Fig. 3C).

**Figure 3.**
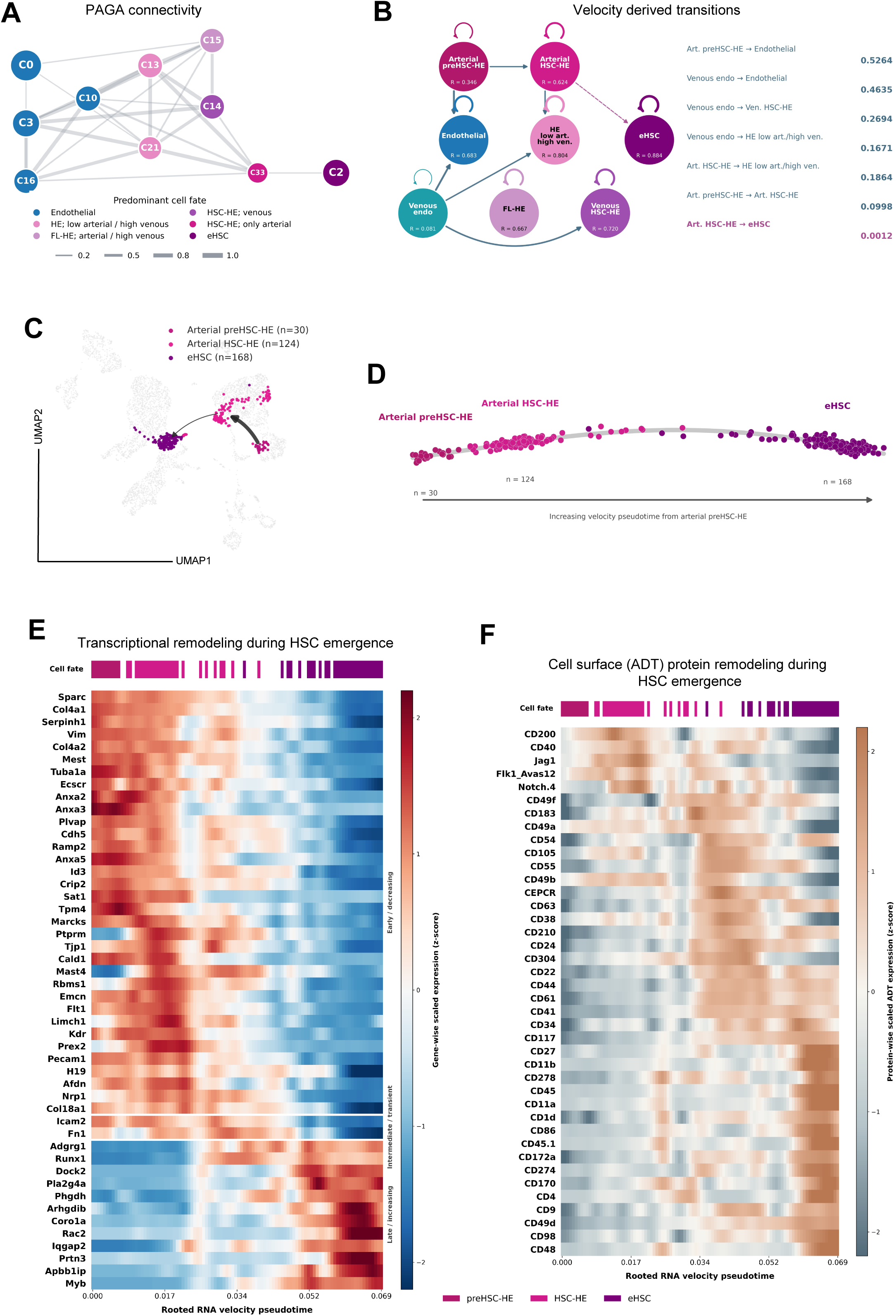
Integrated connectivity and pseudotime analyses identify transcriptional and phenotypic changes associated with HSC emergence. **(A)** Partition-based graph abstraction (PAGA) showing connectivity among selected endothelial, hemogenic endothelial and HSC-associated clusters. Nodes represent annotated clusters, with colours indicating predominant cell identities. Edge thickness and intensity represent relative graph connectivity. (B) RNA velocity-derived transition network among selected endothelial, arterial preHSC-HE, arterial HSC-HE, venous-associated hemogenic and emerging HSC (eHSC) populations. Arrows indicate inferred directional relationships, and the accompanying values summarize velocity-derived transition weights. (C) UMAP highlighting arterial preHSC-HE (n = 30), arterial HSC-HE (n = 124) and eHSC (n = 168) populations within the integrated atlas. Arrows illustrate the inferred direction of progression between these states. **(D)** Rooted RNA velocity pseudotime ordering of arterial preHSC-HE, arterial HSC-HE and eHSC populations. Individual cells are positioned according to their inferred pseudotemporal coordinates and coloured by their annotated identities. The trajectory is rooted in an E9.5 arterial preHSC-HE cell. **(E)** Heatmap showing transcriptional remodelling across the rooted RNA velocity pseudotime trajectory. Selected genes display changing expression patterns between arterial preHSC-HE, arterial HSC-HE and eHSC populations. Columns are ordered by pseudotime, and expression values are standardized by gene. **(F)** Heatmap showing corresponding changes in ADT-derived cell-surface protein abundance along the same rooted trajectory. Columns are ordered by pseudotime, and values are standardized by protein. The upper annotation bars in (E) and (F) indicate the assigned cell identities.

Together, these findings revealed considerable transcriptional heterogeneity among endothelial and HSC-associated populations and identified differences in their inferred developmental relationships and temporal representation. Rather than supporting a single homogeneous transition from endothelium towards HSC-associated identity, these analyses suggested a more complex organization of endothelial and hematopoietic states during embryonic development. Notably, among the endothelial and HE-associated populations examined, arterial HSC-HE (C33) provided the principal graph-based connection to the C2/eHSC population (Fig. 3A).

To further investigate these inferred developmental relationships, we analysed RNA velocity-derived transition probabilities (Fig. 3B–D and Supplementary Fig. 3D). The resulting transition network identified predicted relationships between endothelial, arterial preHSC-HE, arterial HSC-HE and other HE-associated populations (Fig. 3B). Importantly, this analysis suggested a potential progression involving early arterial endothelial populations, arterial preHSC-HE and arterial HSC-HE towards emerging HSC-associated states. To investigate these relationships in greater detail, we examined the RNA velocity-derived connectivity among the three HSC-associated populations (Fig. 3C,D). Arterial preHSC-HE cells displayed predicted connectivity towards arterial HSC-HE, which in turn showed relationships with C2/eHSC (Fig. 3B,C). We further examined this organization using rooted RNA velocity pseudotime, setting arterial preHSC-HE as the initial population and using the velocity-derived transition structure to order cells along the inferred trajectory. This analysis placed arterial HSC-HE between arterial preHSC-HE and eHSC, supporting a transcriptional continuum linking these populations (Fig. 3D).

Additionally, we implemented a third, complementary computational approach to examine the relative positioning of these populations within the broader endothelial- to-hematopoietic landscape. We applied diffusion pseudotime^31^ (DPT) analysis to order individual cells along an inferred developmental continuum. DPT estimates the relative position of cells from a defined starting population based on their connectivity within the transcriptional landscape, providing a continuous representation of potential developmental relationships. We therefore examined the distribution of endothelial, hematopoietic progenitor and eHSC populations along the pseudotime axis (Supplementary Fig. 3E). As expected, endothelial cells generally occupied earlier pseudotemporal positions, whereas hematopoietic stem and progenitor populations were represented towards later positions. Notably, HSC-associated populations did not occupy uniform pseudotemporal positions, suggesting that transcriptionally related cells represent distinct molecular states within the broader endothelial-to-hematopoietic landscape (Supplementary Fig. 3E). Together with the PAGA and RNA velocity analyses, these findings were consistent with an inferred progression from arterial preHSC-HE through arterial HSC-HE towards eHSC-associated identity, although the direction and functional outcome of these relationships remain to be experimentally established.

Since the acquisition of HSC-associated identity is accompanied by extensive molecular changes, we next investigated the transcriptional programmes and cell-surface protein remodelling associated with this inferred progression. Rather than performing a conventional differential gene expression analysis between discrete populations, we examined gene expression patterns along the ordered continuum from arterial preHSC-HE through arterial HSC-HE to eHSC (Fig. 3E). Several genes associated with endothelial and early hemogenic identity showed higher expression towards earlier positions, whereas other genes became enriched towards later positions, consistent with progressive remodelling of transcriptional programmes during the acquisition of HSC-associated identity (Fig. 3E).

Importantly, analysis of antibody-derived surface-protein abundance across the same ordered populations revealed corresponding changes in immunophenotypic characteristics (Fig. 3F). Individual surface proteins displayed distinct patterns of abundance across the inferred developmental continuum, with FLK1 and JAG1 enriched towards earlier endothelial and preHSC-associated states, whereas CD34, CD27, c-KIT (CD117), CD45 and CD274 (PD-L1) became more prominent within later HSC-associated populations (Fig. 3F). These observations demonstrate that the transcriptional differences separating these populations are accompanied by changes in their surface-protein profiles, providing an opportunity to identify phenotypic features associated with distinct molecular states during HSC emergence.

Together, these analyses reveal substantial molecular and immunophenotypic heterogeneity among arterial hemogenic populations and support an inferred transcriptional progression from arterial preHSC-HE through arterial HSC-HE towards emerging HSC-associated identity.

### Distinct arterial hemogenic populations reveal a developmental progression towards HSC-associated identity

Having identified endothelial heterogeneity associated with the emergence of HE states, we next investigated the relationship between arterial preHSC-HE, arterial HSC-HE and emerging HSC-associated populations. These populations occupied related but distinguishable regions of the integrated transcriptional landscape (Fig. 4A), and RNA velocity-derived connectivity suggested potential developmental relationships between them (Fig. 3B–D). In particular, RNA velocity and diffusion pseudotime (DPT) analyses supported an inferred progression from arterial preHSC-HE through arterial HSC-HE towards eHSC (Fig. 3C,D and Supplementary Fig. 3E). Remarkably, the arterial preHSC-HE population was embedded within the broader C0 endothelial cluster (Fig. 1Ci). Moreover, arterial preHSC-HE cells were predominantly detected at E9.5 (Fig. 4Bi,ii and Supplementary Fig. 4A), whereas arterial HSC-HE cells were represented across later developmental stages (Fig. 4Bi,ii). Although both populations were present within the early embryonic compartment, their relative representation changed markedly during development. At E9.5, arterial preHSC-HE constituted the majority of cells within the combined arterial preHSC-HE and arterial HSC-HE compartment, whereas arterial HSC-HE became the predominant population at E10.25 and E11.5 in the AGM (Fig. 4Bi,ii). These findings indicate that arterial preHSC-HE represents an earlier transcriptional state than arterial HSC-HE and are consistent with a potential developmental relationship between these populations.

**Figure 4.**
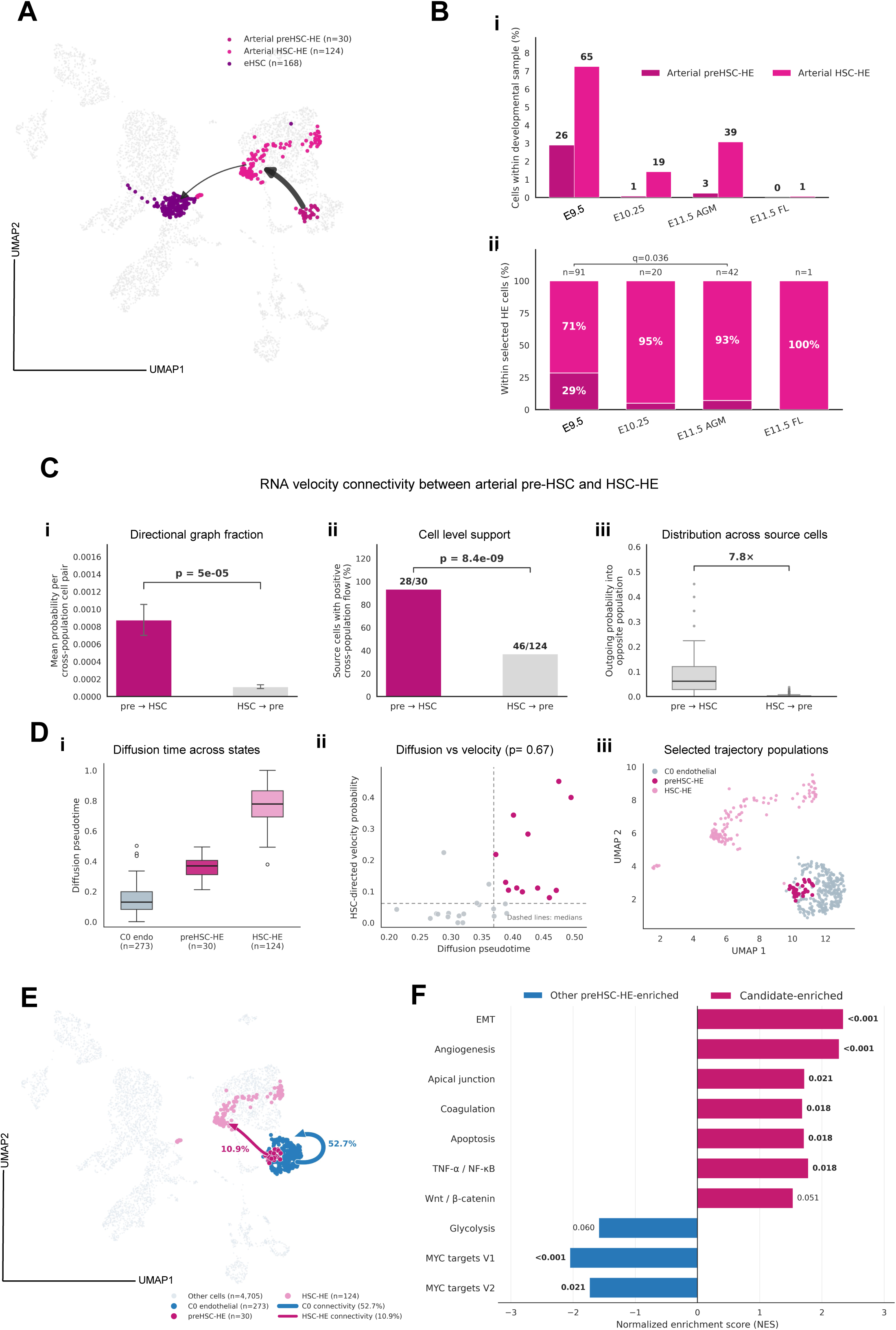
Directional RNA velocity connectivity distinguishes arterial preHSC-HE and HSC-HE populations. **(A)** UMAP highlighting arterial preHSC-HE (n = 30), arterial HSC-HE (n = 124) and eHSC (n = 168) populations within the integrated dataset. Arrows indicate inferred relationships among these transcriptional states. **(B)** Developmental distribution of arterial preHSC-HE and arterial HSC-HE populations. **(i)** Percentage of cells assigned to each population across E9.5, E10.25, E11.5 AGM and E11.5 foetal liver samples, with cell counts indicated above the bars. **(ii)** Relative composition of the combined arterial preHSC-HE and arterial HSC-HE compartment at each developmental stage. The indicated statistical comparison is based on cell-level observations. **(C)** RNA velocity-derived directional connectivity between arterial preHSC-HE and arterial HSC-HE. **(i)** Comparison of the mean directional graph fraction between preHSC-HE-to-HSC-HE and reverse connectivity. **(ii)** Relationship between diffusion pseudotime and HSC-directed RNA velocity connectivity among selected cells. The association was assessed using Spearman’s rank correlation (ρ = 0.67). **(iii)** Distribution of outgoing connectivity values across individual source cells, showing a 7.8-fold difference in the reported directional connectivity measure. Statistical comparisons are indicated in the panels. **(D)** Diffusion pseudotime and RNA velocity analyses of endothelial and arterial hemogenic populations. **(i)** Distribution of diffusion pseudotime values for C0 endothelial cells (n = 273), arterial preHSC-HE (n = 30) and arterial HSC-HE (n = 124). **(ii)** Relationship between diffusion pseudotime and HSC-directed RNA velocity connectivity among selected cells. **(iii)** UMAP showing the selected endothelial, arterial preHSC-HE and arterial HSC-HE populations used for the analysis. **(E)** UMAP representation of RNA velocity-derived connectivity from arterial preHSC-HE towards selected destination populations. The indicated weighted graph fractions include connectivity towards C0 endothelial cells (52.7%) and arterial HSC-HE (10.9%). These values represent inferred connectivity weights rather than experimentally observed cell-fate outcomes. **(F)** Hallmark gene set enrichment analysis comparing arterial preHSC-HE cells with elevated predicted connectivity towards HSC-HE against the remaining arterial preHSC-HE cells. Positive normalized enrichment scores indicate pathways enriched in the candidate population, whereas negative scores indicate enrichment in the comparison population. FDR-adjusted values are displayed alongside the pathways.

To further investigate this relationship, we examined RNA velocity-derived transition probabilities between arterial preHSC-HE and arterial HSC-HE populations (Fig. 4Ci–iii). The inferred connectivity was stronger in the direction from arterial preHSC-HE towards arterial HSC-HE than in the reciprocal direction (Fig. 4Ci,ii). Furthermore, a higher proportion of arterial preHSC-HE cells exhibited detectable predicted connectivity towards arterial HSC-HE than was observed in the reverse direction (Fig. 4Ciii). These findings suggest that the transcriptional dynamics of arterial preHSC-HE are preferentially aligned with those of arterial HSC-HE, supporting a directional relationship between these molecular states.

We next compared the relative pseudotemporal positions of endothelial, arterial preHSC-HE and arterial HSC-HE populations (Fig. 4Di). Endothelial cells predominantly occupied earlier positions along the inferred trajectory, followed by arterial preHSC-HE and arterial HSC-HE populations (Fig. 4Di). To investigate whether individual cells differed in their inferred developmental dynamics, we compared diffusion pseudotime with RNA velocity-derived connectivity towards HSC-associated states (Fig. 4Dii). This analysis revealed a positive association between pseudotemporal position and predicted connectivity (Spearman’s ρ = 0.67), together with considerable heterogeneity among individual cells. Notably, only a subset of cells exhibited elevated predicted connectivity, suggesting that cells within the same transcriptionally defined population may differ in their inferred relationships with downstream HSC-associated states (Fig. 4Dii).

To determine whether these populations could be distinguished within the broader integrated dataset, we examined the distribution of endothelial, arterial preHSC-HE and arterial HSC-HE populations across the UMAP (Fig. 4Diii). This analysis highlighted a restricted population of cells associated with the inferred progression towards HSC-associated identity within the broader endothelial compartment, including the C0 cluster (Fig. 4Diii). These observations further support the presence of transcriptionally distinguishable states within the developing endothelial and hemogenic compartments. Importantly, the velocity-derived connectivity analysis indicated that only a limited component of the inferred transition structure linked the early endothelial compartment towards arterial preHSC-HE, whereas stronger connectivity was associated with endothelial states (Fig. 5A,B). These values represent computationally inferred relationships rather than the proportion of cells that will undergo hemogenic specification or acquire HSC identity.

**Figure 5.**
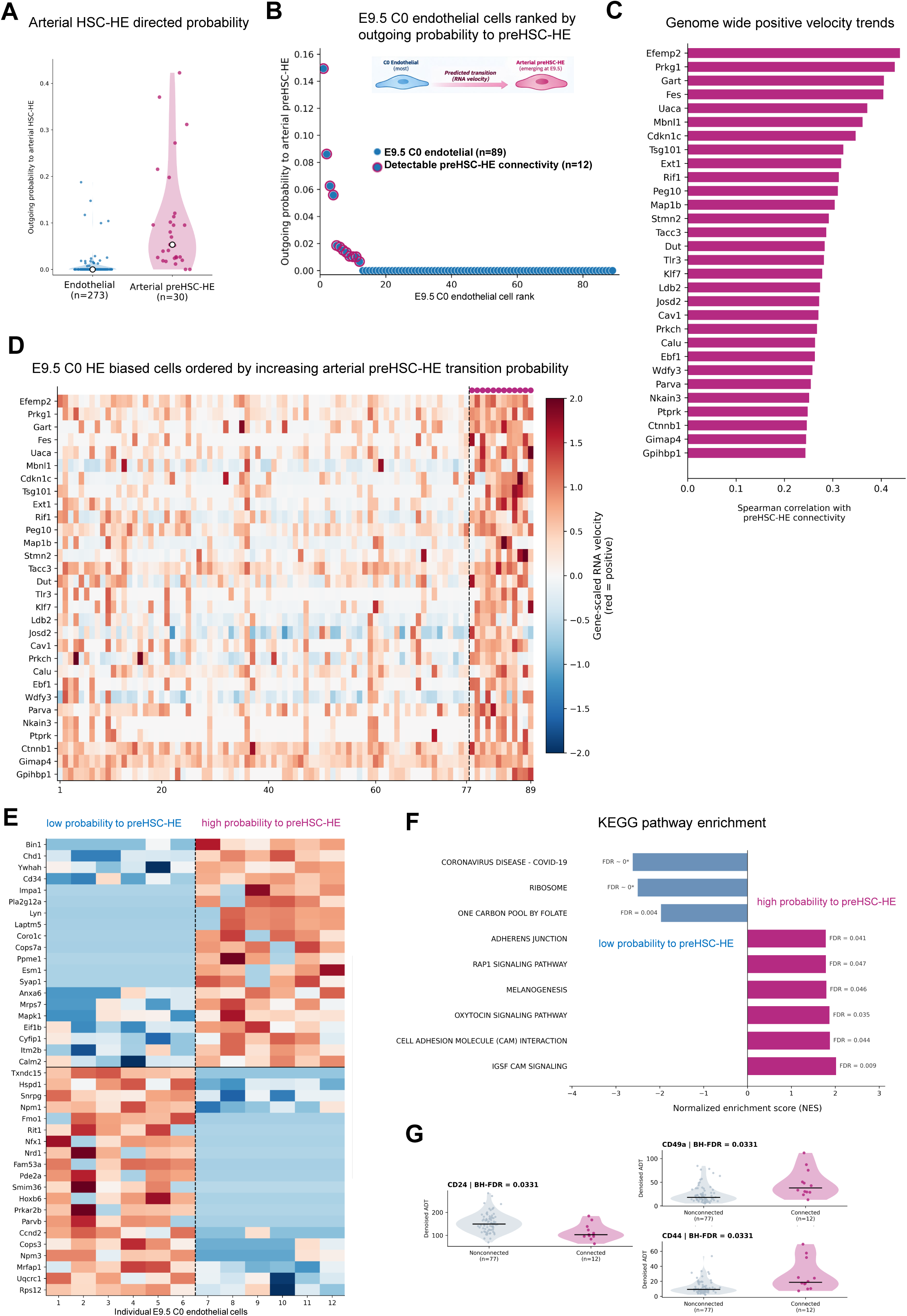
A rare E9.5 endothelial population exhibits increased predicted connectivity towards arterial preHSC-HE and distinct molecular features. **(A)** Distribution of RNA velocity-derived connectivity towards arterial HSC-HE among C0 endothelial cells (n = 273) and arterial preHSC-HE cells (n = 30). Individual points represent cells, and violin plots summarize the distributions. **(B)** E9.5 C0 endothelial cells (n = 101) ranked according to their outgoing RNA velocity-derived connectivity towards arterial preHSC-HE. Cells with detectable positive connectivity (n = 12) are distinguished from those without detectable connectivity (n = 89). **(C)** Genes exhibiting positive associations between expression and predicted connectivity towards arterial preHSC-HE among E9.5 C0 endothelial cells. Bars represent Spearman correlation coefficients. **(D)** Heatmap showing expression of connectivity-associated genes across E9.5 C0 endothelial cells ordered by increasing predicted connectivity towards arterial preHSC-HE. The subset with detectable connectivity is indicated. Values represent standardized gene-expression measurements. **(E)** Heatmap of selected differentially expressed genes comparing E9.5 C0 endothelial cells with low or undetectable connectivity and those with elevated connectivity towards arterial preHSC-HE. Columns represent individual cells, and rows represent genes. Expression values are standardized by gene. **(F)** KEGG pathway enrichment analysis comparing endothelial populations defined by predicted connectivity towards arterial preHSC-HE. Positive normalized enrichment scores indicate pathways enriched in the high-connectivity group, whereas negative scores indicate enrichment in the low-connectivity group. FDR-adjusted values are displayed **(G)** Violin plots comparing ADT-derived CD24, CD49a and CD44 surface-protein abundance between E9.5 C0 endothelial cells with and without detectable predicted connectivity towards arterial preHSC-HE. Individual points represent cells, and FDR-adjusted values are indicated.

Together, these findings reveal distinct arterial hemogenic populations that differ in their developmental representation, transcriptional identity and RNA velocity-derived connectivity. The combined analyses support a model in which arterial preHSC-HE represents an early HSC-associated state with predicted relationships towards arterial HSC-HE, consistent with a potential progression towards emerging HSC identity.

We further investigated the biological programmes associated with these candidate populations using pathway enrichment analysis (GESA) (Fig. 5F). Candidate cells exhibited enrichment of pathways commonly associated with HSC emergence, including epithelial-to-mesenchymal transition ^32–34^, angiogenesis ^9^, apical junction organization ^35,36^, coagulation, apoptosis, TNFα/NF-κB ^37,38^ signalling and WNT/β-catenin signalling ^18,39^. In contrast, other endothelial populations showed relative enrichment of distinct metabolic programmes (Fig. 5F). These findings suggest that the transition towards HSC-associated identity is accompanied by changes in signalling, cellular interactions and transcriptional programmes associated with endothelial remodelling and hematopoietic specification.

### A transcriptionally distinct E9.5 endothelial population exhibits predicted connectivity towards arterial preHSC-HE

The observation that arterial preHSC-HE cells were embedded within the broader endothelial C0 cluster prompted us to investigate whether we could identify endothelial cells associated with the emergence of this HSC-related hemogenic population. Although the developmental origin of hemogenic endothelium has been extensively investigated, the molecular characteristics of endothelial states preceding HSC-associated hemogenic specification remain incompletely understood. To address this question, we examined RNA velocity-derived connectivity from C0 endothelial cells towards arterial preHSC-HE, using the annotated arterial preHSC-HE population as a reference (Fig. 5A). Remarkably, a small subset of E9.5 C0 endothelial cells (12 of 101) exhibited elevated predicted connectivity towards arterial preHSC-HE compared with the remaining endothelial population (Fig. 5B). Importantly, these cells lacked detectable *Runx1* expression, suggesting that they had not acquired a detectable *Runx1*-associated hemogenic transcriptional programme. Although *Procr* expression was elevated in endothelial cells with predicted connectivity towards preHSC-HE, *Mecom* expression was lower than in the comparison population, providing no clear evidence of established HE identity (Supplementary Fig. 4B). These observations suggested that the E9.5 endothelial compartment contains molecularly distinct cells with an inferred relationship to emerging HSC-associated hemogenic populations.

We next examined the predicted destinations of individual E9.5 C0 endothelial cells using one-step RNA velocity-derived transition probabilities (Fig. 5B and Supplementary Fig. 4C). Although predicted connectivity towards endothelial states predominated in most cells, a restricted subset exhibited increased connectivity towards arterial preHSC-HE and other HSC-associated hemogenic states. We subsequently referred to the endothelial cells displaying elevated predicted connectivity towards arterial preHSC-HE as C0 HE-biased cells. This designation reflects their computationally inferred transcriptional relationships rather than experimentally demonstrated hemogenic potential.

To identify the molecular characteristics of C0 HE-biased cells, we examined genes whose expression correlated with predicted connectivity towards arterial preHSC-HE (Fig. 5C,D). This analysis identified a group of genes positively associated with elevated transition probabilities, including *Efemp2*, *Plxg1*, *Gart*, *Fes*, *Uaca*, *Mbnl1*, *Cdh1c*, *Tsg101* and *Ext1*. We further examined the expression of these genes across individual E9.5 endothelial cells ordered according to their predicted transition probabilities (Fig. 5D). Cells with the highest predicted connectivity exhibited increased expression of several genes identified in the correlation analysis, revealing a distinct transcriptional profile within a subset of the endothelial population. These findings suggest that endothelial cells exhibiting predicted relationships with arterial preHSC-HE can be distinguished molecularly before the acquisition of a broader hemogenic transcriptional programme.

To investigate this population in greater detail, we compared the transcriptional profiles of C0 HE-biased and C0 non-biased endothelial cells using differential gene expression analysis (Fig. 5E). This analysis identified transcriptional differences between the two populations, and subsequent gene set enrichment analysis revealed distinct biological pathways associated with their molecular profiles (Fig. 5F). Among the pathways enriched in C0 HE-biased cells were adherens junction, Rap1 signalling, melanogenesis, oxytocin signalling and cell adhesion-related pathways. In contrast, other pathways showed relative enrichment in the remaining endothelial population (Fig. 5F). These observations suggest that differences in cell adhesion, intercellular interactions and signalling may be associated with endothelial transcriptional states exhibiting predicted connectivity towards arterial preHSC-HE. Interestingly, several of the identified genes and pathways differed from the established transcriptional programmes commonly associated with HSC emergence, suggesting that these cells represent a previously unresolved endothelial state rather than a population already displaying a conventional HSC-associated transcriptional identity.

We next examined whether the transcriptional differences between these endothelial populations were accompanied by changes in surface-protein abundance (Fig. 5G). Comparison of antibody-derived tag (ADT) abundance identified differences in CD24, CD49a and CD44 between C0 HE-biased and C0 non-biased endothelial cells. These findings suggest that the transcriptional heterogeneity associated with predicted hemogenic specification is also reflected in the immunophenotypic characteristics of individual endothelial cells.

Together, these analyses identify a molecularly and phenotypically distinct subset of early embryonic endothelial cells associated with increased RNA velocity-derived connectivity towards arterial preHSC-HE. Although these computationally inferred relationships do not establish direct developmental ancestry, they suggest that the early endothelial compartment contains heterogeneous molecular states that may precede the emergence of HSC-associated hemogenic populations.

## Discussion

HSC-generating potential is restricted to a rare subset of hemogenic endothelial cells, whereas a broader population gives rise to hematopoietic progenitors with limited or absent long-term self-renewal capacity^7,9,15,40^. Distinguishing these populations remains challenging because conventional surface-marker combinations do not fully resolve their underlying transcriptional identities. By integrating single-cell transcriptomic and surface-protein measurements, our study provides a multimodal view of embryonic hematopoiesis, revealing substantial heterogeneity among endothelial and hemogenic populations and identifying molecular and phenotypic states associated with HSC emergence.

A central finding is that hemogenic specification and the acquisition of HSC-associated identity are molecularly distinguishable. Although endothelial, arterial, venous and HE/EHT transcriptional programmes identified overlapping populations, HSC-associated signatures were concentrated within a more restricted compartment. Importantly, even endothelial cells expressing *Runx1* but lacking detectable CD41 surface protein displayed heterogeneous hematopoietic lineage-associated transcriptional programmes. These observations suggest that molecular differences associated with hematopoietic output may already be detectable within early hemogenic populations, before overt hematopoietic differentiation. This is consistent with evidence that HSCs and hematopoietic progenitors can emerge through distinct developmental pathways ^7,9^ and supports the view that hemogenic endothelium comprises multiple molecular states rather than a uniform population progressing towards a single hematopoietic fate.

An important advantage of our approach is the ability to directly relate transcriptionally defined populations to their surface-protein profiles. Although combinations of surface markers have long been used to enrich emerging HSCs and hematopoietic progenitors, the cellular identities captured by individual markers remain incompletely characterized. Our integrated analysis reveals that established markers are distributed across multiple transcriptionally distinct populations, highlighting the limitations of interpreting immunophenotype independently of molecular identity. Conversely, the correspondence between *Hlf* expression, HSC-associated transcriptional programmes and CD27 surface abundance identifies CD27 as a potentially useful marker for enriching HSC-associated populations. This relationship is particularly relevant because HLF-expressing embryonic precursors can contribute to both HSC and progenitor populations^7,18^. Thus, while CD27 may help distinguish cells exhibiting an HSC-associated molecular programme, neither CD27 nor *Hlf* expression alone establishes functional HSC identity. More broadly, these findings demonstrate how paired RNA and surface-protein profiling can refine the interpretation of established immunophenotypic markers and inform the design of prospective isolation strategies.

Our analyses also reveal distinct arterial hemogenic states associated with HSC development. Graph-based connectivity, RNA velocity and pseudotime analyses identified relationships among arterial preHSC-HE, arterial HSC-HE and emerging HSC-associated populations. Arterial preHSC-HE was predominantly represented at E9.5, whereas arterial HSC-HE became more prominent at later developmental stages. Importantly, RNA velocity connectivity was strongly asymmetric between these populations, with 28 of 30 arterial preHSC-HE cells showing positive connectivity towards arterial HSC-HE, compared with 46 of 124 cells in the reverse direction. Together with their relative pseudotemporal positions, these observations support an inferred progression from arterial preHSC-HE towards HSC-associated states. These inferred developmental relationship between these populations were accompanied by changes in transcriptional programmes and surface-protein expression. Earlier states were enriched for FLK1 and JAG1, whereas CD34, CD27, c-KIT, CD45 and PD-L1 became more prominent in later HSC-associated populations. The early presence of JAG1 is particularly interesting given the stage- and context-dependent functions of NOTCH signalling during embryonic hematopoiesis ^40–42^. Altogether, these observations are particularly relevant to the longstanding challenge of distinguishing successive stages of HSC development using surface markers. Rather than defining a single universal HSC phenotype, our results suggest that marker expression should be interpreted in the context of developmental stage and underlying transcriptional state. The combined RNA-protein measurements therefore provide a basis for identifying marker combinations that distinguish closely related populations along the inferred HSC-associated developmental continuum.

Perhaps the most intriguing finding is the identification of a rare subset of E9.5 C0 endothelial cells exhibiting elevated RNA velocity-derived connectivity towards arterial preHSC-HE. These cells were distinguishable from the remaining endothelial population despite lacking clear transcriptional evidence of established hemogenic commitment, including detectable *Runx1* expression. Their distinct gene-expression profiles and enrichment of pathways associated with cell adhesion and intercellular signalling suggest that molecular heterogeneity relevant to hemogenic specification may arise before the activation of conventional HE programmes. This raises the possibility that endothelial cells differ in their propensity to acquire HSC-associated hemogenic characteristics before they become recognizable as hemogenic endothelium. However, the observed connectivity does not establish that these cells are committed precursors or that they necessarily progress towards HSC formation.

Importantly, the candidate endothelial population also exhibited differences in surface-protein abundance, providing an opportunity to connect its predicted developmental relationship with an experimentally accessible phenotype. This is particularly valuable because identifying early endothelial populations associated with HSC emergence remains difficult using established HE and hematopoietic markers alone. The combination of transcriptional features, velocity-derived connectivity and surface-protein differences offers a potential strategy for enriching these cells and testing their developmental potential. Prospective isolation followed by functional assays will be essential to determine whether they exhibit increased hemogenic activity, preferentially generate HSC-associated progeny or instead represent a transient endothelial state without a distinct developmental fate.

Together, our findings position HSC-associated development within a heterogeneous and dynamically changing endothelial landscape. By identifying an early endothelial population with predicted connectivity towards preHSC-HE and linking distinct transcriptional states to accessible surface markers, this study provides a framework for testing how HSC-associated identity emerges. Prospective isolation, functional assays and lineage tracing will be essential to establish whether these molecular states represent committed HSC precursors or transient windows of developmental competence.

## Materials and Methods

### Mouse embryo generation and processing

Mouse work was performed in accordance with the United Kingdom Animal Scientific Procedures Act (ASPA) 1986. Animal experiments performed at the Cancer Research United Kingdom Manchester Institute (CRUK-MI) were approved by the Animal Welfare and Ethics Review Body of the CRUK-MI. Experiments performed at the University of Oxford were approved by the Oxford Clinical Medicine Ethical Review Committee. Mice were housed in individually ventilated cages under standard conditions, including a 12-h light-dark cycle, ambient temperature of 19-23 °C and relative humidity of 45-65%, in accordance with UK Home Office guidelines and institutional protocols. The wild type mouse lines CD1 we used to generate embryos used in this study. Vaginal plug detection was considered as E0.5 and staging was confirmed for each embryo at the time of collection by visual inspection.

For the single cell CiteSeq AGMs of E9.5 (n=9), E10.25 (n=23) and E11.5 (n=10) embryos were dissected in PBS with 7% foetal calf serum (FBS) and penicillin/streptomycin (100 U/mL). Single-cell suspensions were generated by incubating the tissues for 20-30 min in 500 ul of 1 mg/ml of Collagenase/Dispase (Roche cat# 10269638001) before mechanical dissociation with a syringe and needle. The resulting single-cell suspension was used to stain with fluorescently labelled antibodies, CD31 and cKIT, together with the panel of ADT conjugated TotalSeqB antibodies staining for 30 min at 4 degrees. Samples were sorted with a FACS AriaII. FACS plots were generated using FlowJo V10.

### Downstream Processing & Sequencing Workflow

TotalSeq-B antibody-stained cells were counted using an Invitrogen Countess III cell counter with Trypan Blue. They were then loaded into a GEM-X OCM chip (10X Genomics, PN-1000747), configured to multiplex 8 samples into 2 pools of 4 samples each. cDNA was prepared using the Chromium Next GEM Single Cell 3’ v4 OCM kit with feature barcoding (10X Genomics, PN-1000779, PN-1000702) according to manufacturer’s instructions. A gene expression library and TotalSeq-B library were prepared for each pool according to the 10X Genomics protocol. Libraries were quality checked using the Agilent Fragment Analyzer and quantified by qPCR using a KAPA Library Quantification Kit for Illumina Platforms (Roche, 07960336001). Sequencing was performed on an Illumina NovaSeq 6000.

### Computational processing of single-cell RNA and surface-protein data

#### Data import, sample identification and cell exclusion

Single-cell RNA sequencing (scRNA-seq) and antibody-derived tag (ADT) data were analyzed from four mouse embryonic sample groups: E9.5, E10.25, E11.5 aorta-gonad-mesonephros (AGM), and E11.5 foetal liver (FL). Count matrices were imported into Python using Scanpy and the read_10x_mtx function with gex_only=False to retain both gene-expression and antibody-capture features. The initial dataset comprised 5,450 cells: 970 E9.5 cells, 1,430 E10.25 cells, 1,342 E11.5 AGM cells, and 1,708 E11.5 FL cells. A predefined list of 313 barcodes was excluded, comprising 72, 90, 72, and 79 cells from the respective sample groups. Barcode identifiers were normalized before matching to the exclusion lists. The correspondence between exclusion-file labels and physical sample folders was established through barcode matching and retained in the exclusion audit. After exclusion, 5,137 cells remained. Samples were concatenated using Scanpy with an outer feature join. Cell identifiers were made unique by appending sample identifiers, and the physical source-sample identity was retained in cell metadata. The resulting multimodal dataset contained 33,696 gene-expression features and 120 antibody-capture features. RNA and ADT measurements were separated into their respective matrices for downstream multimodal analysis.

#### totalVI integration of RNA and ADT measurements

RNA and cell-surface protein measurements were integrated using totalVI, implemented in scvi-tools (version 1.3.3). The model was configured using RNA counts stored in the counts layer and ADT measurements stored in protein_expression. Sample identity was included as a batch covariate. Of the 120 ADT features, two were excluded from the model-training copy because they contained zero counts across all cells (NOTCH3 and E-Cadherin). The original analysis object retained all 120 features, while totalVI was trained using 118 ADT markers. A totalVI model with 20 latent dimensions was initialized using random seed 42. Training was performed on an NVIDIA L4 GPU, with a maximum of 200 epochs, early stopping enabled, and validation checks every five epochs. The trained model was saved, and the resulting 20-dimensional latent representation was extracted for all 5,137 cells. This representation was stored in X_totalVI and used for neighborhood construction, clustering, and visualization.

#### Neighborhood graph construction, clustering and UMAP

A nearest-neighbor graph was constructed using the totalVI latent representation, with 15 neighbors, Euclidean distance, and random seed 0. Leiden clustering was performed at resolution 2.4 using the leidenalg implementation, two iterations, and random seed 42. This analysis identified 39 clusters, retained in the annotated dataset. A two-dimensional UMAP embedding was generated for visualization of the integrated multimodal landscape. The saved UMAP metadata specify parameters *a* = 0.5830300203414425 and *b* = 1.3341669924314914, with random seed 42. The totalVI embedding and corresponding cell annotations provided the reference framework for subsequent analyses of endothelial, hemogenic endothelial, and hematopoietic populations.

#### Cell-fate annotation and gene-program analysis

Cell populations were annotated using their integrated transcriptional profiles, lineage-associated markers, and cell-surface protein expression. The annotation framework distinguished endothelial populations, arterial and venous-associated endothelial and hemogenic endothelial states, arterial preHSC-HE, arterial HSC-HE, emerging HSCs (eHSCs), and differentiated hematopoietic populations. Transcriptional programs associated with endothelial identity, arterial specification, venous specification, hemogenic endothelial/endothelial-to-hematopoietic transition (HE/EHT), and HSC-associated identity were evaluated using Scanpy’s score_genes function. For program scoring, RNA counts were normalized to 10,000 counts per cell and log-transformed. Program scores were calculated using random seed 42 and visualized across the totalVI-derived UMAP.

#### Developmental composition and hematopoietic subpopulation analysis

Cell populations were grouped according to developmental sample and final cell-fate annotation. Population abundances were summarized using cell counts and relative frequencies within the relevant sample or parent compartment. The distributions of arterial preHSC-HE and arterial HSC-HE were compared across developmental stages. Selected hematopoietic populations, including lymphoid-primed progenitors, were also examined for developmental changes in abundance. Where indicated, categorical associations were evaluated using contingency-table procedures and Fisher’s exact tests, with Benjamini-Hochberg correction for multiple comparisons.

#### RNA velocity preprocessing and estimation

RNA velocity was analyzed using scVelo (version 0.3.4). The multimodal and velocity datasets were matched using cell identifiers, yielding 5,132 matched cells from the 5,137-cell multimodal dataset. Genes were retained for velocity analysis when total spliced counts exceeded three and total unspliced counts exceeded three. This filtering retained 15,972 genes from the original 33,696 RNA features. The existing neighborhood graph based on the totalVI representation was reused for the calculation of spliced and unspliced neighborhood moments. RNA velocity was estimated using the stochastic model. The saved velocity parameters were mode=“stochastic”, mode_neighbors=“distances”, n_recurse_neighbors=2, perc=[5,95], and fit_offset=False. The resulting velocity object retained spliced and unspliced counts, neighborhood moments (Ms and Mu), estimated velocities, and positive and negative velocity graphs. The full velocity graph had dimensions 5,132 × 5,132.

#### RNA velocity connectivity between annotated populations

The RNA velocity graph was used to investigate predicted directional relationships among endothelial, hemogenic endothelial, and emerging hematopoietic populations. Cell-level velocity connectivity was summarized for selected source and destination annotations. These analyses were used to visualize directional relationships, quantify outgoing connectivity, and compare the support for transitions between arterial preHSC-HE and arterial HSC-HE. The analysis distinguished positive connectivity toward a destination population from the absence of detectable positive connectivity. For arterial preHSC-HE and arterial HSC-HE, positive forward connectivity was detected for 28 of 30 arterial preHSC-HE cells, compared with positive reverse connectivity for 46 of 124 arterial HSC-HE cells. The recorded source-cell-normalized directional graph fraction was approximately 7.8-fold higher in the preHSC-HE-to-HSC-HE direction. Additional analyses compared arterial preHSC-HE connectivity with endothelial and HSC-HE populations. The reported weighted connectivity fractions toward C0 endothelial cells and arterial HSC-HE were 52.7% and 10.9%, respectively.

#### Diffusion pseudotime and connectivity-associated endothelial states

Diffusion pseudotime was used to examine transcriptional ordering among selected endothelial and hemogenic endothelial populations. Analyses included C0 endothelial cells, arterial preHSC-HE, and arterial HSC-HE. Pseudotime distributions were compared across annotated populations. Associations between pseudotime and velocity-derived connectivity were evaluated using Spearman rank correlation where specified in the analysis. Endothelial cells were additionally ranked by their inferred connectivity toward arterial preHSC-HE. Gene-expression patterns associated with this ranking were examined using correlation and differential-expression approaches. A comparism between high-connectivity and other endothelial cells was performed using Mann-Whitney U tests. RNA expression was normalized to 10,000 counts per cell and log-transformed. Resulting *P* values were adjusted using the Benjamini-Hochberg procedure. In that implementation, genes were selected using FDR < 0.05 and an absolute log2 fold-change threshold of at least 0.5. Selected endothelial-associated surface proteins were visualized using ADT violin plots.

#### Gene set enrichment analysis (GSEA)

Gene set enrichment analysis was used to investigate transcriptional programs associated with velocity-defined endothelial and hemogenic endothelial subpopulations. For the arterial preHSC-HE analysis, 30 cells were separated into 12 HSC-HE-biased candidates and 18 comparison cells. An E9.5-restricted sensitivity analysis compared eight HSC-HE-biased candidates with 18 other arterial preHSC-HE cells. We used two preranked GSEA implementations using GSEApy and MSigDB Hallmark gene sets. Genes were ranked using a signed expression-effect score standardized by a variance estimate with regularization. Genes expressed in fewer than three cells across the compared groups were excluded from ranking. GSEA was performed using 10,000 permutations, gene-set size limits of 5-500, and random seed 42. Normalized enrichment scores, nominal *P* values, and false discovery rate-adjusted values were retained. For the pathway-analysis workflow, we used KEGG enrichment results, with the final plotting code retaining pathways meeting FDR < 0.05.

#### Rooted RNA velocity pseudotime

A focused RNA velocity trajectory was constructed from arterial preHSC-HE, arterial HSC-HE, and eHSC populations. The selected dataset contained 322 cells, comprising 30 arterial preHSC-HE cells, 124 arterial HSC-HE cells, and 168 eHSCs.

Velocity information was derived from the full 5,132-cell graph, and an E9.5 arterial preHSC-HE cell (CTTCTATGTGATAGTG-1-E9) was selected as the root for the focused pseudotime analysis. The resulting median rooted velocity pseudotime values were 0.001887 for arterial preHSC-HE, 0.015702 for arterial HSC-HE, and 0.062022 for eHSC. The corresponding pseudotime ranges were 0-0.008388, 0.008398-0.039366, and 0.025680-0.068757, respectively. Cells were ordered according to rooted velocity pseudotime for visualization and subsequent analyses of molecular remodeling. Pseudotime was interpreted as a computational ordering rather than chronological developmental time.

#### Identification of E9.5 C0 endothelial cells with predicted connectivity towards arterial preHSC-HE

To identify endothelial cells exhibiting predicted connectivity towards arterial preHSC-HE, we selected cells annotated as C0 endothelial cells from the E9.5 sample (n = 101). Using the RNA velocity graph generated with scVelo under the stochastic model, we examined the outgoing velocity-derived connectivity from each endothelial cell towards cells annotated as arterial preHSC-HE. For each source cell, positive connectivity weights directed towards the arterial preHSC-HE population were used to distinguish cells with detectable predicted connectivity from those without detectable connectivity. This analysis identified 12 endothelial cells with positive connectivity towards arterial preHSC-HE, whereas the remaining 89 cells showed no detectable positive connectivity towards this population. These two groups were subsequently used to investigate connectivity-associated transcriptional differences, gene-expression correlations, pathway enrichment and surface-protein abundance. The classification was based exclusively on RNA velocity-derived connectivity and did not involve selection using established hemogenic markers such as *Runx1*.

#### Modeling transcriptional dynamics along rooted velocity pseudotime

Transcriptional dynamics were analyzed across the 322-cell rooted velocity trajectory. RNA counts were normalized to 10,000 counts per cell and transformed using log1p. Genes were required to be detected in at least 17 cells, corresponding to the greater of 10 cells or 5% of the trajectory population. For each retained gene, expression was modeled as a function of normalized rooted pseudotime using a cubic B-spline basis with three internal knots positioned at pseudotime quantiles. The fitted spline model was compared with an intercept-only model using an F-test. Nominal *P* values were adjusted using the Benjamini-Hochberg procedure. A total of 11,212 genes were evaluated in the saved trajectory statistics. Dynamic genes were selected using FDR < 0.05 and a fitted expression range of at least 0.25. The retained dynamic-gene results contained 6,120 genes.The final transcriptional heatmap displayed 48 selected genes. Cells were ordered by rooted pseudotime, and expression was summarized over 100 pseudotime bins, with interpolation across unoccupied bins. Gaussian smoothing was applied, and expression values were standardized by gene for visualization. The final heatmap implementation used Gaussian smoothing with σ = 4 and clipped standardized values to the interval −2.5 to 2.5. Gene order and transcriptional-pattern classifications were retained in the accompanying heatmap-order table.

#### Modeling cell-surface protein dynamics along rooted velocity pseudotime

ADT dynamics were analyzed for cells matched to the rooted RNA velocity trajectory. The analysis retrieved protein-expression measurements from the saved RNA-ADT AnnData object and aligned them with the cell-level rooted pseudotime values. The implementation checked the numerical characteristics of the ADT matrix and applied a log1p transformation when measurements were identified as strongly count-like. Protein abundance was modeled as a function of normalized pseudotime using cubic polynomial regression. The polynomial model was compared with an intercept-only model using an F-test, followed by Benjamini-Hochberg correction. A total of 120 proteins were evaluated, and 40 proteins were selected for the final trajectory heatmap using adjusted significance and fitted-expression-range criteria. Protein expression was summarized across 100 pseudotime bins, interpolated across missing bins, and smoothed using a Gaussian filter with σ = 2. Protein-wise standardized values were displayed along the common rooted pseudotime coordinate. Protein labels were simplified for visualization by removing technical antibody-species prefixes where appropriate.

#### Statistical analysis and interpretation

Statistical procedures included Mann-Whitney U tests, Fisher’s exact tests, Spearman rank correlation, regression-model F-tests, and preranked gene set enrichment analysis, according to the specific analysis. Multiple-testing correction was performed using the Benjamini-Hochberg method where specified in the code. Nominal significance values are reported as *P* values and adjusted values as FDR or *q* values.

#### Software and reproducibility

Analyses were performed in Python using Google Colab. Additional software used across the notebooks included Scanpy, AnnData, NumPy, SciPy, pandas, PyTorch, Matplotlib, Seaborn, scikit-learn, UMAP-learn, GSEApy, and statsmodels. The saved analysis files include the annotated totalVI dataset, the stochastic RNA velocity dataset, the full RNA velocity graph, the trained totalVI model checkpoint, and the rooted trajectory statistics. Additionally, reproducibility was independently verified by the Computational Biology support group, CRUK as following:

Bioinformatic Validation: Clustering & Cell Annotation Cite-seq data containing the RNA and ADT assays were loaded into R (v 4.6.1) using the Seurat package (v5.6.0)1. The RNA assay was filtered to remove cells < 200 features and > 8000 features. The data was also filtered to exclude dying cells using a mitochondrial percentage threshold of > 5. This removed 313 suspected dying cells, reducing total cell count from 5,449 to 5,136. Cells were clustered using the Louvain algorithm using the top 16 principal components (PCs) and with a resolution of 1, producing 25 clusters. Differential expression analysis was performed using the FindAllMarkers function and genes were filtered (padj<= 0.05). All significant DEGs per cluster (ordered by log2FC) were exported to “RNA_SignificantDEGs_Annotated.csv” and filtered by log2FC >=1 to “RNA_SignificantDEGs_Annotated_Log2FC_Filtered.csv”. To annotate higher resolution cell types required for downstream analyses, HSC and endothelial clusters were selected for re-clustering. Weighted Nearest Neighbour Analysis (WNN) was performed using the FindMultiModalNeighbors function to cluster the cells using both the RNA and ADT modalities. The top 20 PCs for each modality and a resolution of 2 was used. RNA weight per annotated cluster was plotted using the VlnPlot function. Differential expression analysis was performed using the FindAllMarkers function and ADTs were filtered (padj<= 0.05). All significant ADTs per cluster (ordered by log2FC) were exported to “ADT_SignificantDEGs_Annotated.csv” and filtered by log2FC >=1 to “ADT_SignificantDEGs_Annotated_Log2FC_Filtered.csv”. Specific panels of ADTs for each cluster were plotted on heatmaps using the DoHeatmap function. RNA Velocity Analysis Loom files containing unspliced and spliced RNA matrices were generated using the Velocyto package (v0.17.17) with the velocyto run command for each of the timepoints2. An in-house R script was used to read-in, convert, and reformat .loom files for each timepoint into a merged .rds object. The barcodes were filtered to include only those which had passed quality checking in the annotated Seurat object. Spliced, un-spliced and raw count RNA matrices, alongside metadata and UMAP coordinates were exported for analysis using the Python package scVelo (v0.3.4)3. An anndata object was created using raw RNA counts, raw spliced/unspliced count matrices, and metadata containing cell type annotations. The object was normalised using the scv.pp.filter_and_normalize function. The scv.pp.moments function was used to neighbourhood-smooth the spliced and un-spliced RNA counts by averaging the expression profiles of each cell with its spatial neighbours in Principal Component Analysis (PCA) space. The scv.pl.proportions function was used to visualise the proportion of spliced vs unspliced RNA across annotated cell types. scv.recover_dynamics was used to fit a gene-specific dynamical model, jointly estimating the transcription/splicing/degradation kinetics and each cell’s position along each gene’s fitted trajectory. scv.tl.velocity was next used to fit each cell’s transcriptional state using the fitted model to infer the speed and direction of expression changes, then scv.tl.velocity_graph was used to derive cell-cell directional relationships, and to infer the directionality of each cell based on its transcriptional changes relative to neighbouring cells.

## Data availability

Single cell RNA-Seq and combined TotalSeqB dataset is available upon request to corresponding authors.

## Acknowledgements

The authors thank the following facilities of the Cancer Research UK Manchester Institute for technical support: Mouse facility, Flow Cytometry, and the Molecular Biology Core Facility. Roshana Thambyrajah is supported by a Ramon y Cajal 2023 award. The Lacaud group is supported by Cancer Research UK, C5759/A20971.

## Contributions

RT and conceptualized the study, designed the experiments, analyzed data and wrote the manuscript. RT, VS, AGG and CSC performed experiments and analyzed data. RT, KS, DGS and SS analyzed the CiteSeq data. SS and RT supervised the CiteSeq data analysis. GL, RT, ML, and FPC supervised data analysis. RT, GL, and ML wrote the manuscript.

## Supplementary figure legends

**Supplementary Figure 1. Experimental cell enrichment and developmental distribution of endothelial and hematopoietic populations in the CITE-seq atlas. (A)** Representative flow cytometry plots illustrating the enrichment of endothelial, hemogenic endothelial (HE) and intra-aortic hematopoietic cluster (IAHC) populations based on CD31 and c-KIT expression. Samples were collected from E9.5 aorta-gonad-mesonephros (AGM; 9 embryos from one plug), E10.25 AGM (23 embryos from two plugs), E11.5 AGM (10 embryos from one plug) and E11.5 foetal liver (FL). Approximately 2,500 endothelial/HE and IAHC cells were targeted for collection from each sample. **(B)** Uniform manifold approximation and projection (UMAP) plots showing the distribution of transcriptionally annotated cell populations across developmental stages. Cells from E9.5 (n = 898), E10.25 (n = 1,340), E11.5 AGM (n = 1,270) and E11.5 foetal liver (n = 1,629) are displayed separately using the integrated totalVI embedding. Cells are coloured according to their assigned cell identities, illustrating developmental and anatomical differences in the representation of endothelial, hemogenic endothelial and hematopoietic populations.

**Supplementary Figure 2. Comparison of transcript expression and surface-protein abundance across the embryonic hematopoietic landscape. (A)** UMAP feature plots comparing RNA expression and totalVI-denoised antibody-derived tag (ADT) abundance for selected cell-surface markers, including CD44 (*Cd44*), EPCR (*Procr*), CD150 (*Slamf1*) and CD27 (*Cd27*). Transcript expression and corresponding surface-protein abundance are displayed on the same integrated UMAP coordinates. RNA expression is shown in red and ADT abundance in brown, with independently scaled colour gradients. **(B)** UMAP feature plots showing the expression of selected transcriptional regulators associated with hemogenic specification and HSC development, including *Runx1*, *Mecom*, *Gata2*, *Mllt3* and *Hoxa9*. Colour intensity represents log-normalized RNA expression.

**Supplementary Figure 3. Heterogeneity of endothelial and hemogenic populations revealed by CD41 expression, lineage-associated transcriptional programmes and computational trajectory analyses. (A)** Distribution of CD41 surface-protein abundance across annotated endothelial and hemogenic endothelial populations. The left panel shows the distribution of denoised CD41 ADT abundance, while the right panel indicates the percentage of cells with undetectable CD41 surface protein within each population. **(B)** Distribution of the highest-scoring hematopoietic lineage-associated transcriptional programme among *Runx1*-expressing endothelial cells lacking detectable CD41 surface protein. Bars indicate the number of cells assigned the highest score for HSC-associated, erythroid, megakaryocytic, myeloid, granulocytic or lymphoid-associated programmes. **(C)** RNA velocity stream plots projected onto the integrated UMAP, illustrating inferred directions of transcriptional change across endothelial, hemogenic endothelial and hematopoietic populations. Velocity vectors were estimated using the stochastic RNA velocity model. **(D)** Additional RNA velocity stream plot showing the inferred transcriptional dynamics across the annotated embryonic hematopoietic landscape. **(E)** Distribution of diffusion pseudotime (DPT) values across selected endothelial, hemogenic endothelial and HSC-associated clusters. Individual cells are coloured according to their annotated cell identities, allowing comparison of their relative positions along the inferred transcriptional continuum.

**Supplementary Figure 4. Arterial preHSC-HE representation and RNA velocity-derived relationships within the early endothelial compartment. (A)** Developmental distribution of arterial preHSC-HE cells within Leiden cluster C0. Stacked bars show the percentage of C0 cells annotated as arterial preHSC-HE and other C0 endothelial cells at E9.5, E10.25, E11.5 AGM and E11.5 foetal liver. Cell counts and percentages are indicated within the bars. **(B)** Comparison of *Runx1*, *Mecom* and *Procr* transcript expression between E9.5 C0 endothelial cells (n = 89) and arterial preHSC-HE cells (n = 26). Individual points represent cells, and the indicated false discovery rate (FDR)-adjusted values summarize the comparisons. **(C)** Stacked bar plots showing one-step RNA velocity-derived transition probabilities for individual E9.5 Leiden C0 cells, including endothelial cells with detectable connectivity towards arterial preHSC-HE (n = 12), endothelial cells without detectable connectivity (n = 77), annotated arterial preHSC-HE cells (n = 26) and other C0 cells (n = 13). Colours indicate predicted destination populations, including endothelial, arterial preHSC-HE, arterial HSC-HE and other annotated states. These values represent computationally inferred transition probabilities rather than experimentally observed developmental outcomes.

