## Supplementary material for "A CITE-seq atlas of mouse embryonic hematopoiesis maps endothelial heterogeneity and reveals emerging HSC-associated states": Suppl Figs

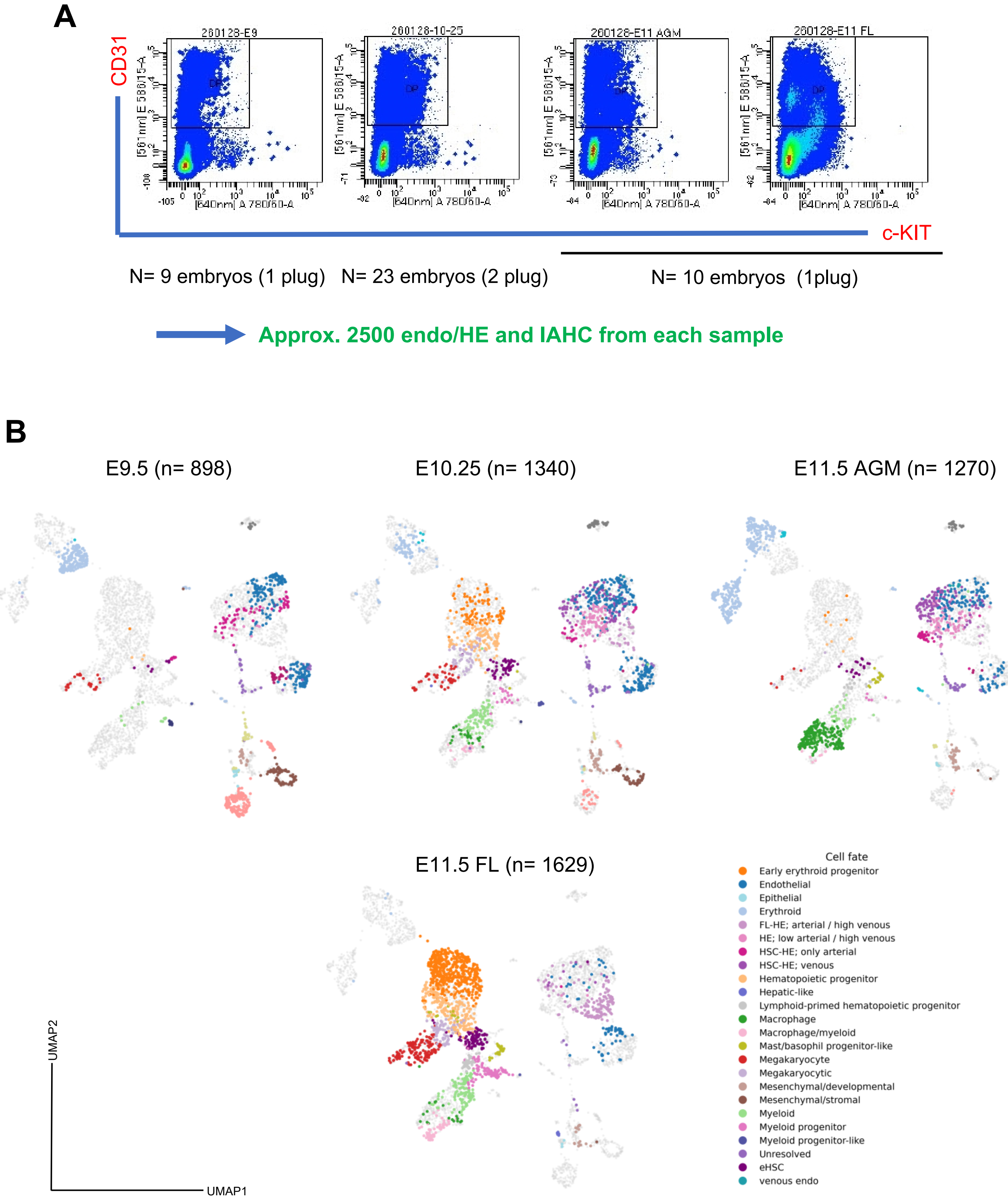

A
CD31
c-KIT
260128-E9
260128-10-25
260128-E11 AGM
260128-E11 FL
DP
N= 9 embryos (1 plug)
N= 23 embryos (2 plug)
N= 10 embryos (1plug)
Approx. 2500 endo/HE and IAHC from each sample
B
E9.5 (n= 898)
E10.25 (n= 1340)
E11.5 AGM (n= 1270)
E11.5 FL (n= 1629)
UMAP2
UMAP1
Cell fate
Early erythroid progenitor
Endothelial
Epithelial
Erythroid
FL-HE; arterial / high venous
HE; low arterial / high venous
HSC-HE; only arterial
HSC-HE; venous
Hematopoietic progenitor
Hepatic-like
Lymphoid-primed hematopoietic progenitor
Macrophage
Macrophage/myeloid
Mast/basophil progenitor-like
Megakaryocyte
Megakaryocytic
Mesenchymal/developmental
Mesenchymal/stromal
Myeloid
Myeloid progenitor
Myeloid progenitor-like
Unresolved
eHSC
venous endo

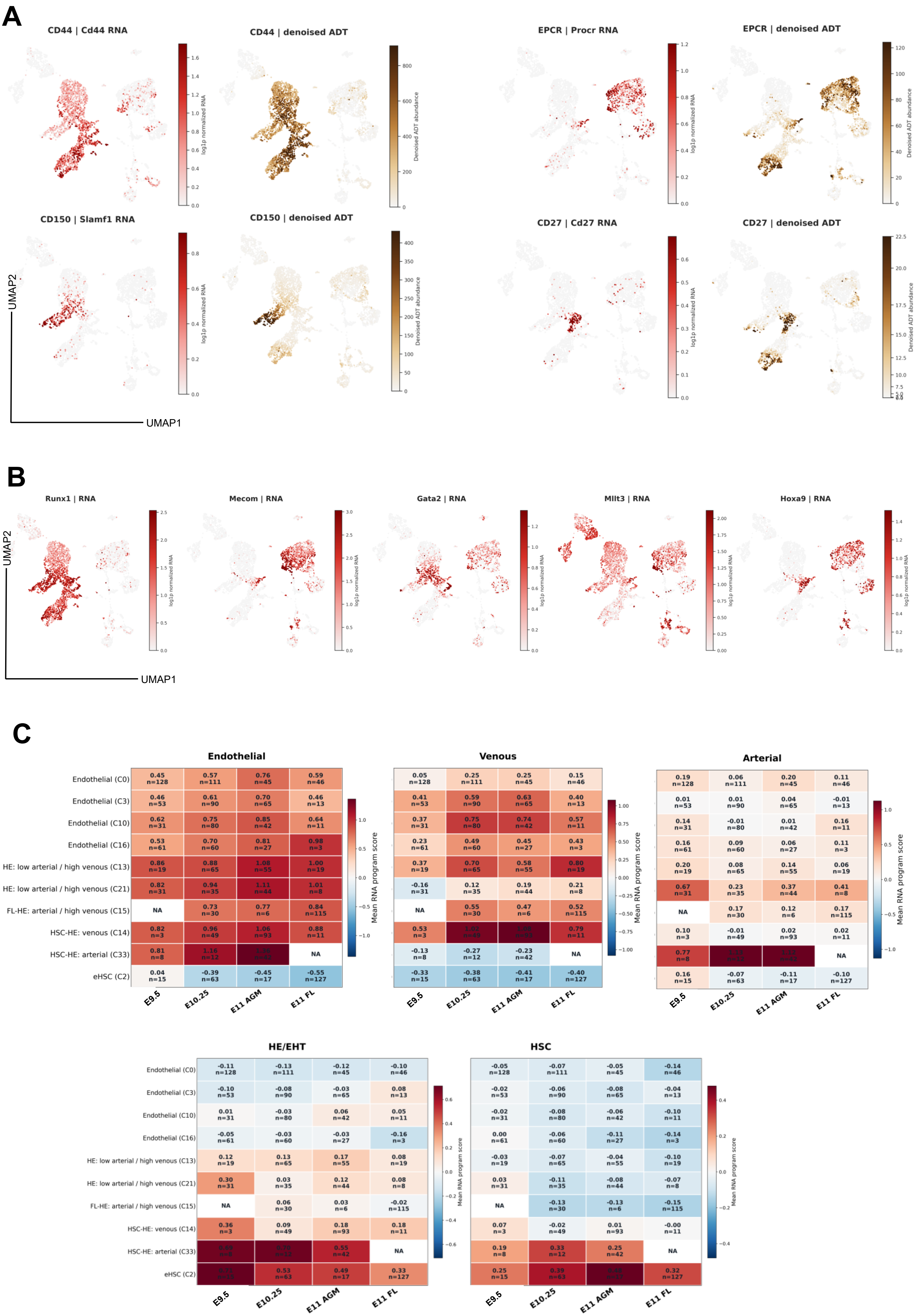
A
CD44 | Cd44 RNA
CD44 | denoised ADT
EPCR | Procr RNA
EPCR | denoised ADT
CD150 | Slamf1 RNA
CD150 | denoised ADT
CD27 | Cd27 RNA
CD27 | denoised ADT
log1p normalized RNA
Denoised ADT abundance
UMAP2
UMAP1
B
Runx1 | RNA
Mecom | RNA
Gata2 | RNA
Mllt3 | RNA
Hoxa9 | RNA
log1p normalized RNA
UMAP2
UMAP1
C
Endothelial
Venous
Arterial
HE/EHT
HSC
Mean RNA program score
Endothelial (C0)
Endothelial (C3)
Endothelial (C10)
Endothelial (C16)
HE: low arterial / high venous (C13)
HE: low arterial / high venous (C21)
FL-HE: arterial / high venous (C15)
HSC-HE: venous (C14)
HSC-HE: arterial (C33)
eHSC (C2)
E9.5
E10.25
E11 AGM
E11 FL
Endothelial: C0: 0.45 n=128, 0.57 n=111, 0.76 n=45, 0.59 n=46
C3: 0.46 n=53, 0.61 n=90, 0.70 n=65, 0.46 n=13
C10: 0.62 n=31, 0.75 n=80, 0.85 n=42, 0.64 n=11
C16: 0.53 n=61, 0.70 n=60, 0.81 n=27, 0.98 n=3
C13: 0.86 n=19, 0.88 n=65, 1.08 n=55, 1.00 n=19
C21: 0.82 n=31, 0.94 n=35, 1.11 n=44, 1.01 n=8
C15: NA, 0.73 n=30, 0.77 n=6, 0.84 n=115
C14: 0.82 n=3, 0.96 n=49, 1.06 n=93, 0.88 n=11
C33: 0.81 n=8, 1.16 n=12, NA
C2: 0.04 n=15, -0.39 n=63, -0.45 n=17, -0.55 n=127
Venous: C0: 0.05 n=128, 0.25 n=111, 0.25 n=45, 0.15 n=46
C3: 0.41 n=53, 0.59 n=90, 0.63 n=65, 0.40 n=13
C10: 0.37 n=31, 0.75 n=80, 0.74 n=42, 0.57 n=11
C16: 0.23 n=61, 0.49 n=60, 0.45 n=27, 0.43 n=3
C13: 0.37 n=19, 0.70 n=65, 0.58 n=55, 0.80 n=19
C21: -0.16 n=31, 0.12 n=35, 0.19 n=44, 0.21 n=8
C15: NA, 0.55 n=30, 0.47 n=6, 0.52 n=115
C14: 0.53 n=3, 0.79 n=11
C33: -0.13 n=8, -0.27 n=12, -0.23 n=42, NA
C2: -0.33 n=15, -0.38 n=63, -0.41 n=17, -0.40 n=127
Arterial: C0: 0.19 n=128, 0.06 n=111, 0.20 n=45, 0.11 n=46
C3: 0.01 n=53, 0.01 n=90, 0.04 n=65, -0.01 n=13
C10: 0.14 n=31, -0.01 n=80, 0.01 n=42, 0.16 n=11
C16: 0.16 n=61, 0.09 n=60, 0.06 n=27, 0.11 n=3
C13: 0.20 n=19, 0.08 n=65, 0.14 n=55, 0.06 n=19
C21: 0.67 n=31, 0.23 n=35, 0.37 n=44, 0.41 n=8
C15: NA, 0.17 n=30, 0.12 n=6, 0.17 n=115
C14: 0.10 n=3, -0.01 n=49, 0.02 n=93, 0.02 n=11
C33: 0.77 n=8, NA
C2: 0.16 n=15, -0.07 n=63, -0.11 n=17, -0.10 n=127
HE/EHT: C0: -0.11 n=128, -0.13 n=111, -0.12 n=45, -0.10 n=46
C3: -0.10 n=53, -0.08 n=90, -0.03 n=65, 0.08 n=13
C10: 0.01 n=31, -0.03 n=80, 0.06 n=42, 0.05 n=11
C16: -0.05 n=61, -0.03 n=60, -0.03 n=27, -0.16 n=3
C13: 0.12 n=19, 0.13 n=65, 0.17 n=55, 0.08 n=19
C21: 0.30 n=31, 0.03 n=35, 0.12 n=44, 0.08 n=8
C15: NA, 0.06 n=30, 0.03 n=6, -0.02 n=115
C14: 0.36 n=3, 0.09 n=49, 0.18 n=93, 0.18 n=11
C33: 0.55 n=42, NA
C2: 0.53 n=63, 0.49 n=17, 0.33 n=127
HSC: C0: -0.05 n=128, -0.07 n=111, -0.05 n=45, -0.14 n=46
C3: -0.02 n=53, -0.06 n=90, -0.08 n=65, -0.04 n=13
C10: -0.02 n=31, -0.08 n=80, -0.06 n=42, -0.10 n=11
C16: 0.00 n=61, -0.06 n=60, -0.11 n=27, -0.14 n=3
C13: -0.03 n=19, -0.07 n=65, -0.04 n=55, -0.10 n=19
C21: 0.03 n=31, -0.11 n=35, -0.08 n=44, -0.07 n=8
C15: NA, -0.13 n=30, -0.13 n=6, -0.15 n=115
C14: 0.07 n=3, -0.02 n=49, 0.01 n=93, -0.00 n=11
C33: 0.19 n=8, 0.33 n=12, 0.25 n=42, NA
C2: 0.25 n=15, 0.39 n=63, 0.32 n=127

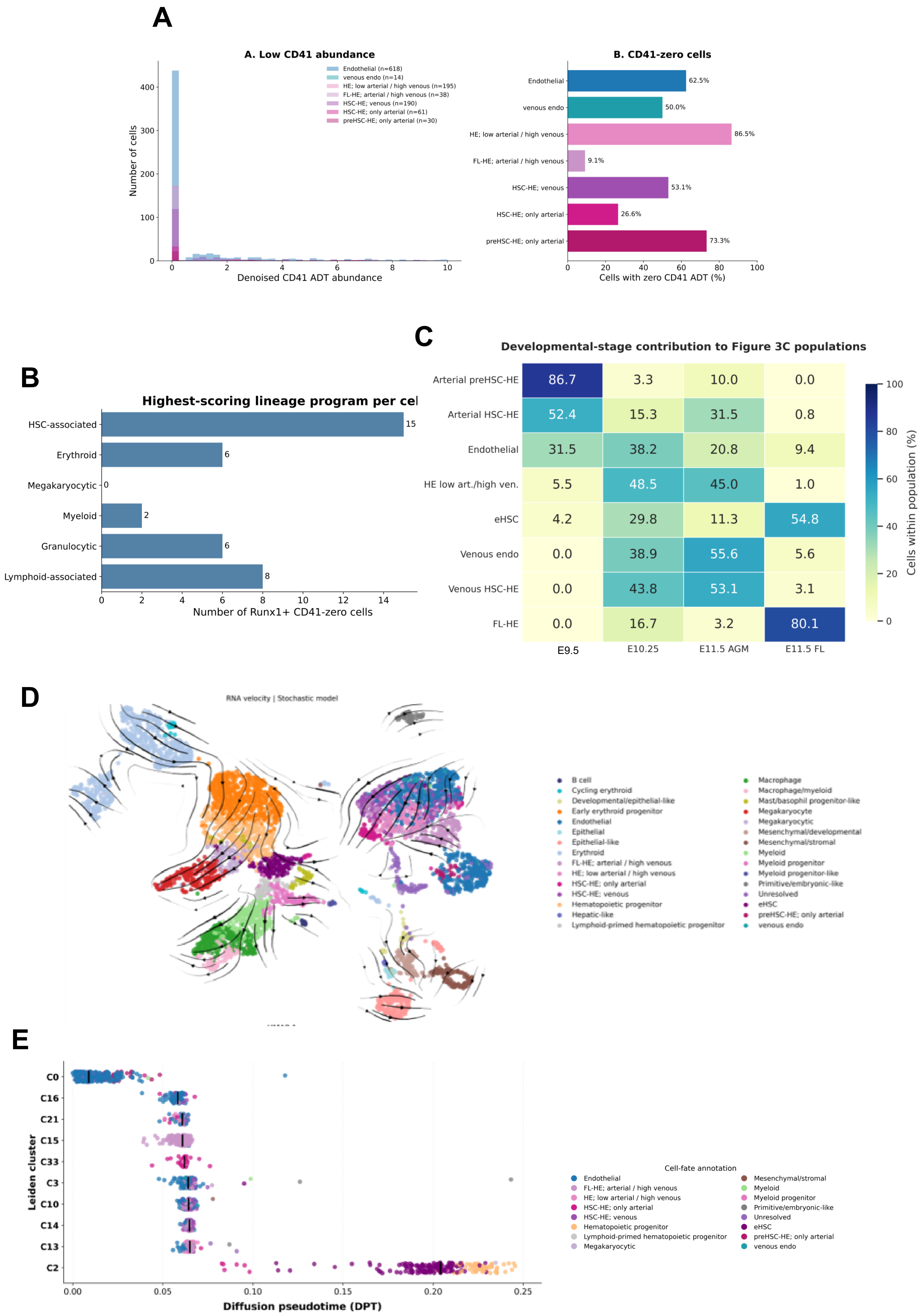

A
A. Low CD41 abundance
Endothelial (n=618)
venous endo (n=14)
HE; low arterial / high venous (n=195)
FL-HE; arterial / high venous (n=38)
HSC-HE; venous (n=190)
HSC-HE; only arterial (n=61)
preHSC-HE; only arterial (n=30)
Number of cells
Denoised CD41 ADT abundance
B. CD41-zero cells
Endothelial 62.5%
venous endo 50.0%
HE; low arterial / high venous 86.5%
FL-HE; arterial / high venous 9.1%
HSC-HE; venous 53.1%
HSC-HE; only arterial 26.6%
preHSC-HE; only arterial 73.3%
Cells with zero CD41 ADT (%)
B
Highest-scoring lineage program per cel
HSC-associated 15
Erythroid 6
Megakaryocytic 0
Myeloid 2
Granulocytic 6
Lymphoid-associated 8
Number of Runx1+ CD41-zero cells
C
Developmental-stage contribution to Figure 3C populations
E9.5
E10.25
E11.5 AGM
E11.5 FL
Arterial preHSC-HE 86.7 3.3 10.0 0.0
Arterial HSC-HE 52.4 15.3 31.5 0.8
Endothelial 31.5 38.2 20.8 9.4
HE low art./high ven. 5.5 48.5 45.0 1.0
eHSC 4.2 29.8 11.3 54.8
Venous endo 0.0 38.9 55.6 5.6
Venous HSC-HE 0.0 43.8 53.1 3.1
FL-HE 0.0 16.7 3.2 80.1
Cells within population (%)
D
RNA velocity | Stochastic model
B cell
Cycling erythroid
Developmental/epithelial-like
Early erythroid progenitor
Endothelial
Epithelial
Epithelial-like
Erythroid
FL-HE; arterial / high venous
HE; low arterial / high venous
HSC-HE; only arterial
HSC-HE; venous
Hematopoietic progenitor
Hepatic-like
Lymphoid-primed hematopoietic progenitor
Macrophage
Macrophage/myeloid
Mast/basophil progenitor-like
Megakaryocyte
Megakaryocytic
Mesenchymal/developmental
Mesenchymal/stromal
Myeloid
Myeloid progenitor
Myeloid progenitor-like
Primitive/embryonic-like
Unresolved
eHSC
preHSC-HE; only arterial
venous endo
E
C0
C16
C21
C15
C33
C3
C10
C14
C13
C2
Leiden cluster
Diffusion pseudotime (DPT)
Cell-fate annotation
Endothelial
FL-HE; arterial / high venous
HE; low arterial / high venous
HSC-HE; only arterial
HSC-HE; venous
Hematopoietic progenitor
Lymphoid-primed hematopoietic progenitor
Megakaryocytic
Mesenchymal/stromal
Myeloid
Myeloid progenitor
Primitive/embryonic-like
Unresolved
eHSC
preHSC-HE; only arterial
venous endo

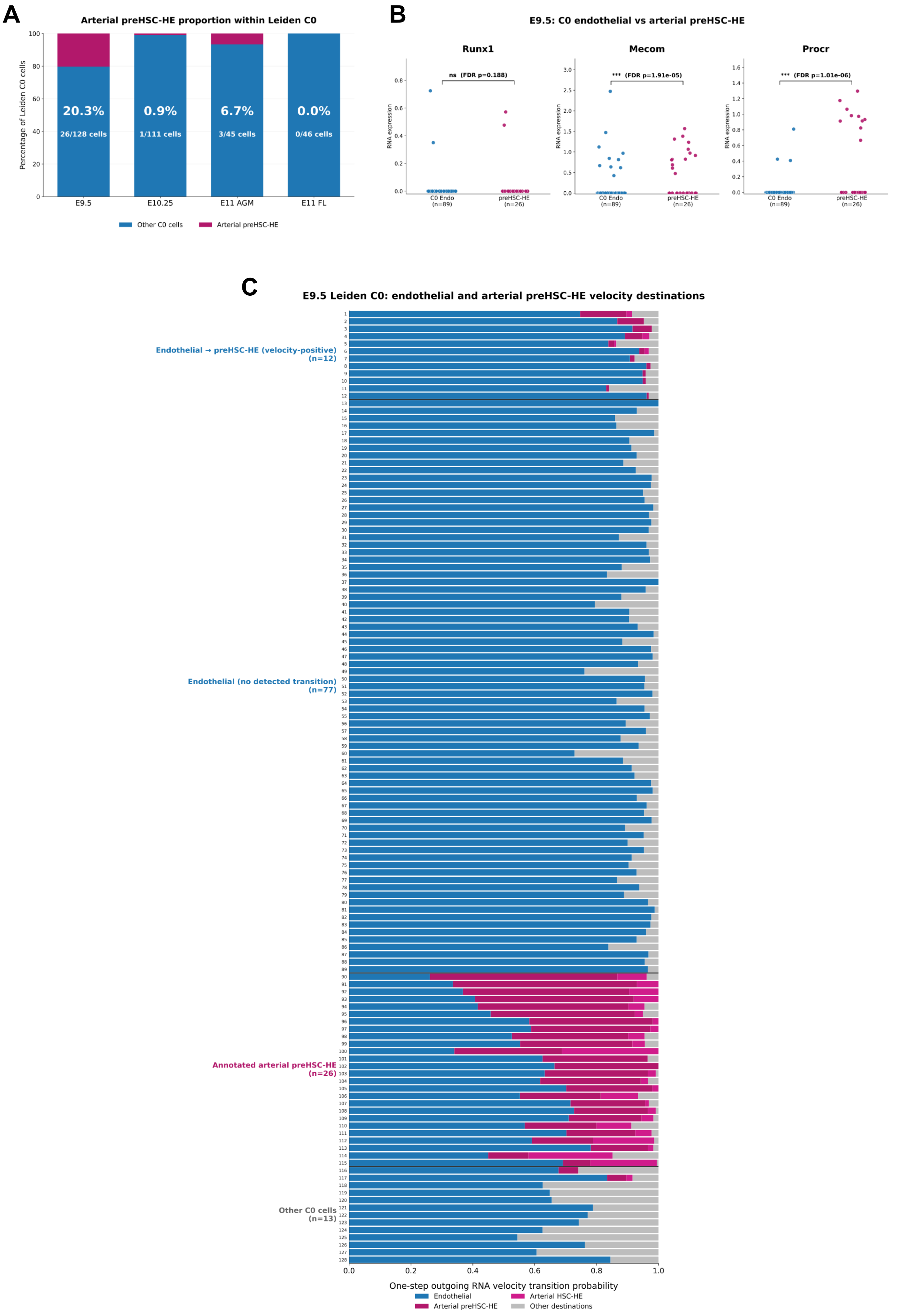

A
Arterial preHSC-HE proportion within Leiden C0
Percentage of Leiden C0 cells
20.3%
26/128 cells
0.9%
1/111 cells
6.7%
3/45 cells
0.0%
0/46 cells
E9.5
E10.25
E11 AGM
E11 FL
Other C0 cells
Arterial preHSC-HE
B
E9.5: C0 endothelial vs arterial preHSC-HE
Runx1
Mecom
Procr
RNA expression
ns (FDR p=0.188)
*** (FDR p=1.91e-05)
*** (FDR p=1.01e-06)
C0 Endo (n=89)
preHSC-HE (n=26)
C
E9.5 Leiden C0: endothelial and arterial preHSC-HE velocity destinations
Endothelial → preHSC-HE (velocity-positive) (n=12)
Endothelial (no detected transition) (n=77)
Annotated arterial preHSC-HE (n=26)
Other C0 cells (n=13)
One-step outgoing RNA velocity transition probability
Endothelial
Arterial preHSC-HE
Arterial HSC-HE
Other destinations
